# Uncovering the Genetic Architecture of Broad Antisocial Behavior through a Genome-Wide Association Study Meta-analysis

**DOI:** 10.1101/2021.10.19.462578

**Authors:** Jorim J. Tielbeek, Emil Uffelmann, Benjamin S. Williams, Lucía Colodro-Conde, Éloi Gagnon, Travis T. Mallard, Brandt Levitt, Philip R. Jansen, Ada Johansson, Hannah Sallis, Giorgio Pistis, Gretchen R.B. Saunders, Andrea G Allegrini, Kaili Rimfeld, Bettina Konte, Marieke Klein, Annette M. Hartmann, Jessica E Salvatore, Ilja M. Nolte, Ditte Demontis, Anni Malmberg, S. Alexandra Burt, Jeanne Savage, Karen Sugden, Richie Poulton, Kathleen Mullan Harris, Scott Vrieze, Matt McGue, William G. Iacono, Nina Roth Mota, Jonathan Mill, Joana F. Viana, Brittany L Mitchell, Jose J Morosoli, Till Andlauer, Isabelle Ouellet-Morin, Richard E. Tremblay, Sylvana Côté, Jean-Philippe Gouin, Mara Brendgen, Ginette Dionne, Frank Vitaro, Michelle K Lupton, Nicholas G Martin, COGA Consortium, Spit for Science Working Group, Enrique Castelao, Katri Räikkönen, Johan Eriksson, Jari Lahti, Catharina A Hartman, Albertine J. Oldehinkel, Harold Snieder, Hexuan Liu, Martin Preisig, Alyce Whipp, Eero Vuoksimaa, Yi Lu, Patrick Jern, Dan Rujescu, Ina Giegling, Teemu Palviainen, Jaakko Kaprio, Kathryn Paige Harden, Marcus R. Munafò, Geneviève Morneau-Vaillancourt, Robert Plomin, Essi Viding, Brian Boutwell, Fazil Aliev, Danielle Dick, Arne Popma, Stephen V Faraone, Anders Børglum, Sarah E Medland, Barbara Franke, Michel Boivin, Jean-Baptiste Pingault, Jeffrey C Glennon, James C. Barnes, Simon E. Fisher, Terrie E. Moffitt, Avshalom Caspi, Tinca JC Polderman, Danielle Posthuma, Broad Antisocial Behavior Consortium collaborators

## Abstract

Despite the substantial heritability of antisocial behavior (ASB), specific genetic variants robustly associated with the trait have not been identified. The present study by the Broad Antisocial Behavior Consortium (BroadABC) meta-analyzed data from 28 discovery samples (N = 85,359) and five independent replication samples (N = 8,058) with genotypic data and broad measures of ASB. We identified the first significant genetic associations with broad ASB, involving common intronic variants in the forkhead box protein P2 *(FOXP2)* gene (lead SNP rs12536335, P = 6.32 x 10^-10^). Furthermore, we observed intronic variation in *Foxp2* and one of its targets *(Cntnap2)* distinguishing a mouse model of pathological aggression (BALB/cJ strain) from controls (BALB/cByJ strain). The SNP-based heritability of ASB was 8.4% (s.e.= 1.2%). Polygenic-risk-score (PRS) analyses in independent samples revealed that the genetic risk for ASB was associated with several antisocial outcomes across the lifespan, including diagnosis of conduct disorder, official criminal convictions, and trajectories of antisocial development. We found substantial genetic correlations of ASB with mental health (depression rg□=□0.63, insomnia rg = 0.47), physical health (overweight rg = 0.19, waist-to-hip ratio rg = 0.32), smoking (rg□=□0.54), cognitive ability (intelligence rg= −0.40), educational attainment (years of schooling rg = −0.46) and reproductive traits (age at first birth rg=□- 0.58, father’s age at death rg= −0.54). Our findings provide a starting point towards identifying critical biosocial risk mechanisms for the development of ASB.

## Main

Antisocial behaviors (ASB) are disruptive acts characterized by covert and overt hostility and violation of the rights and safety of others^1^. The emotional, social, and economic costs incurred by victims of antisocial behavior are far-reaching, ranging from victims’ psychological trauma to reduced productivity when victims miss work to costs incurred by taxpayers in order to staff and run a justice system^2,3^. ASB has been recognized not merely as a social problem, but also as a mental health economic priority^4^. In addition of causing harm to others, those with ASB are themselves at elevated risk of criminal convictions as well as mental health and substance abuse problems^5^. Given all this, it is a research imperative to illuminate the mechanisms underlying the pathogenesis, emergence, and persistence of ASB.

Toward this end, statistical genetic studies have consistently revealed the relevance of environmental and genetic risk factors in the genesis of inter-individual differences in ASB. Family studies – mostly conducted in samples of European ancestry – have demonstrated a considerable heritable component for ASB, with estimates of approximately 50%^6^ across studies. The increasing availability of genomewide data along with data on dimensional ASB measures facilitates in building more advanced explanatory models aimed at identifying trait-relevant genetic variants, that could serve as moderators of socio-environmental factors and vice versa. Moreover, while heritability estimates can differ across subtypes of ASB (e.g., significantly higher twin-based heritability estimates for aggressive forms (65%) versus non-aggressive, rule-breaking forms (48%) of antisocial behavior^7^), these subtypes are genetically correlated (r_g_ = .38)^8^.

### Measuring antisocial behaviour, a broad view

Considering multiple forms of ASB together increases power of genetic analysis and may improve our ability to detect new genetic variants. Here, we thus examine a broadly defined construct of antisocial behaviors, an approach that has successful precedents. Large-scale genomic studies have indicated substantial genetic overlap among psychiatric disorders^9^. A recent genome-wide metaanalysis across eight neuropsychiatric disorders revealed extensive pleiotropic genetic effects (N = 232,964 cases and 494,162 controls)^10,11^. The study found that 109 out of the total 146 contributing loci were associated with at least two psychiatric disorders, suggesting broad liability to these conditions. Moreover, the Externalizing Consortium recently conducted a multivariate analysis of large-scale genome-wide association studies (GWAS) of seven externalizing-related phenotypes (N= ~1.5 million) and found 579 genetic associations with a general liability to externalizing behavior^12^. Although these very large multivariate approaches are crucial in enhancing genetic discovery across phenotypes, they do not detect all the genetic variation relevant to individual disorders. Since ASB is a critical issue for psychiatry and for society, the present study uniquely focuses on (severe) forms of ASB and persistence over the lifespan. To do so, we initiated the Broad Antisocial Behavior Consortium (BroadABC), to perform large-scale meta-analytical genetic analyses, utilizing a broad range of phenotypic ASB measures (e.g., conduct disorder symptoms, aggressive behavior, and delinquency). In our first meta-analysis^13^, we demonstrated that effect sizes for SNPs with suggestive evidence of association with ASB were small, as anticipated for most polygenic traits. Still, we found that the collective effect across all of the included variants (typically referred to as ‘SNP heritability’) explained roughly 5% of the total variation in ASB^13^, which is in line with meta-analyses of the ACTION^14^ and EAGLE^15^ consortium.

To date, however, no previous GWAS meta-analysis targeting broad ASB detected SNPs or genes that are well-replicated. The polygenic architecture of ASB underscores the importance of employing very large samples to yield sufficient power to detect genetic loci of small effect size. Therefore, we substantially boost statistical power by quadrupling the sample size and adding new cohorts to the BroadABC consortium. Since ASB is a critical issue for psychiatry and for society, the present study uniquely focuses on (severe) forms of ASB and persistence over the lifespan.

In our meta-analysis, we also include the results of a GWAS study of Disruptive Behavior Disorders (DBDs) in the context of Attention-Deficit/Hyperactivity Disorder (ADHD), which identified three genome-wide significant loci for DBDs^16^. The present study considers multiple measures of antisocial behaviors in people with and without psychiatric diagnoses across 28 samples to reveal the genetic underpinnings of ASB phenotypes typically studied in psychology, psychiatry, and criminology. These larger samples allow well-powered genetic correlation analyses and improved polygenic risk scores (PRS). Five independent cohorts (total N = 8,058) were employed to validate the ASB PRS in different populations, at different developmental stages, and for different ASB phenotypes. Moreover, we conducted a follow-up analysis of significant loci by using a mouse model of pathological aggression. Since ASB is known to correlate phenotypically with an array of cognitive and health problems^17^, we tested for genetic overlap between ASB and a range of other traits and disorders, including anthropometric, cognitive, reproductive, neuropsychiatric, and smoking.

## Results

### Meta-analysis on broad ASB identifies association with common variants in FOXP2

After quality control and imputation to the Haplotype Reference Consortium or 1000 Genomes Project reference panel (see **Online Methods**), 85,359 individuals from 28 cohorts and a maximum of 7,392,849 variants were available for analysis. We carried out a pooled-sex GWAS meta-analysis for the broad ASB phenotype with METAL^18^ and found one genome-wide significant locus, on chromosome 7 (chromosome band 7q31.1, **Fig. 1A, Supplementary Table 3**). The top lead SNP was rs12536335 (P = 6.32 x 10^-10^; **Fig. 1B and 1C**), located in an intronic region upstream of one of the transcriptional start-sites for the forkhead box protein P2 *(FOXP2)* gene^19,20^. Consistent with this finding, a gene-based association test carried out with MAGMA^21^, identified significant association for *FOXP2 (P =* 7.43 x 10^-7^, **Supplementary Note 3, Supplementary Figure 1, Supplementary Table 6**). The *FOXP2* gene has been related to the development of speech and language^22^, yet is also implicated in a wide range of other traits and diagnoses^23^ (see **Fig. 1D**). MAGMA generalized geneset and tissue-specific gene-set analyses (sex-combined) yielded no significant gene-sets after Bonferroni-correction for multiple testing. The top gene-set for generalized gene-set analysis was activated NTRK2 signals through RAS signaling pathway, **Supplementary Table 7**, while the top tissue-specific gene expression was the hypothalamus, **Supplementary Table 8**). We next ran sexspecific GWAS meta-analyses. These analyses did not identify SNPs that reached genome-wide significance (**Supplementary Tables 4 and 5**).

**Figure 1.**
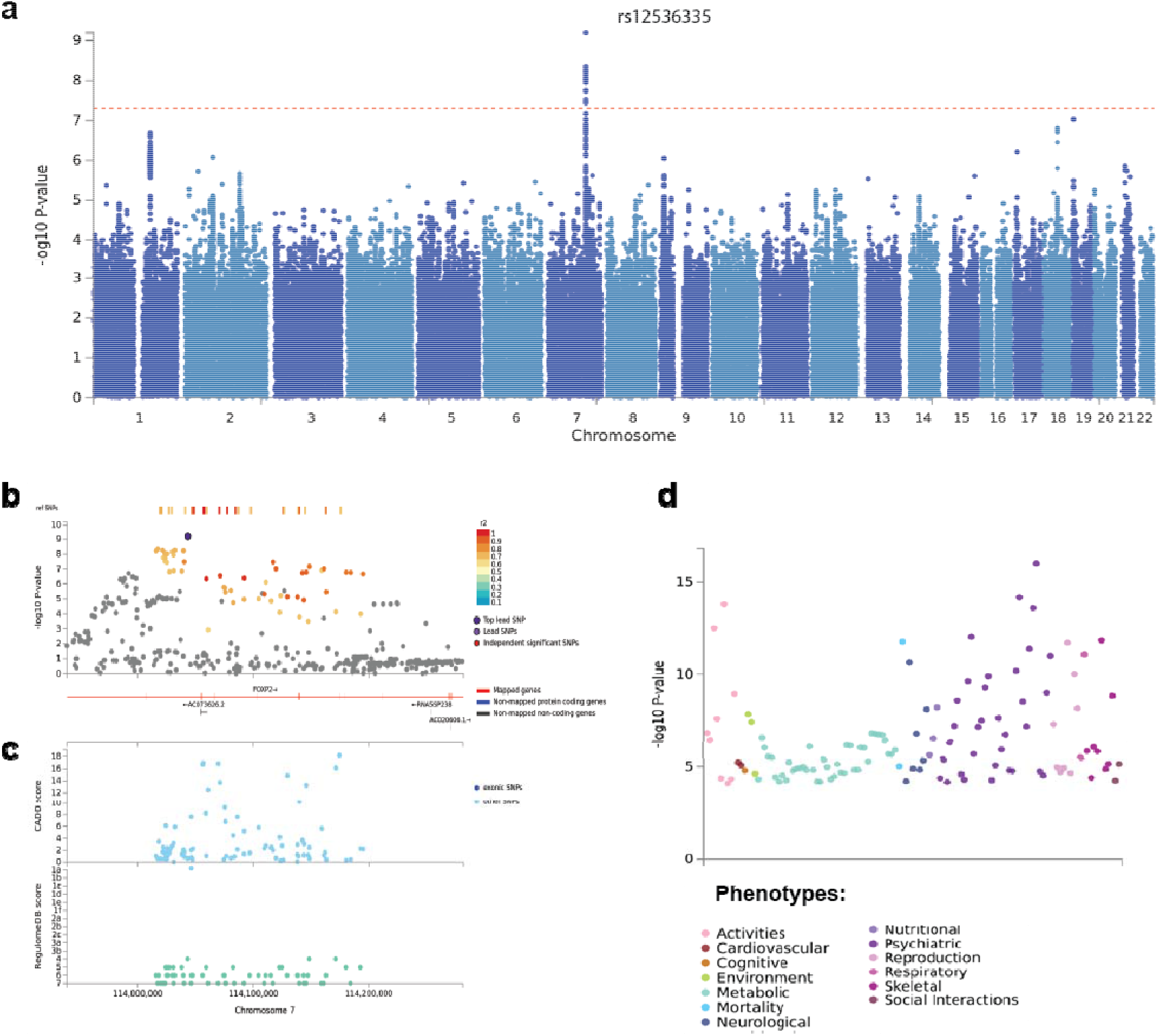
SNP-based results from the GWAS meta-analysis of broad ASB. **A**. Manhattan plot of the GWAS meta-analysis (N = 85,359) of a broad antisocial behavior phenotype, showing the negative log10-transformed P value for each SNP. SNP two-sided P values from a linear model were calculated using METAL^18^, weighting SNP associations by sample size. **B**. Regional association plot around chromosome 7:114043159 with functional annotations of SNPs in LD of lead SNP rs12536335 (shown in purple). The plot displays GWAS P-value plotted against its chromosomal position, where colors represent linkage disequilibrium and r^2^ values with the most significantly associated SNP. **C.** The plot displays CADD scores (Combined Annotation Dependent Depletion) and RegulomeDB scores of these SNPs. **D**. PheWAS plot showing the significance of associations of common variation in the *FOXP2* gene with a wide range of traits and diagnoses based on MAGMA gene-based tests (with Bonferroni corrected P-value: 1.05e-5), as obtained from GWASAtlas (https://atlas.ctglab.nl).

**Figure 2:**
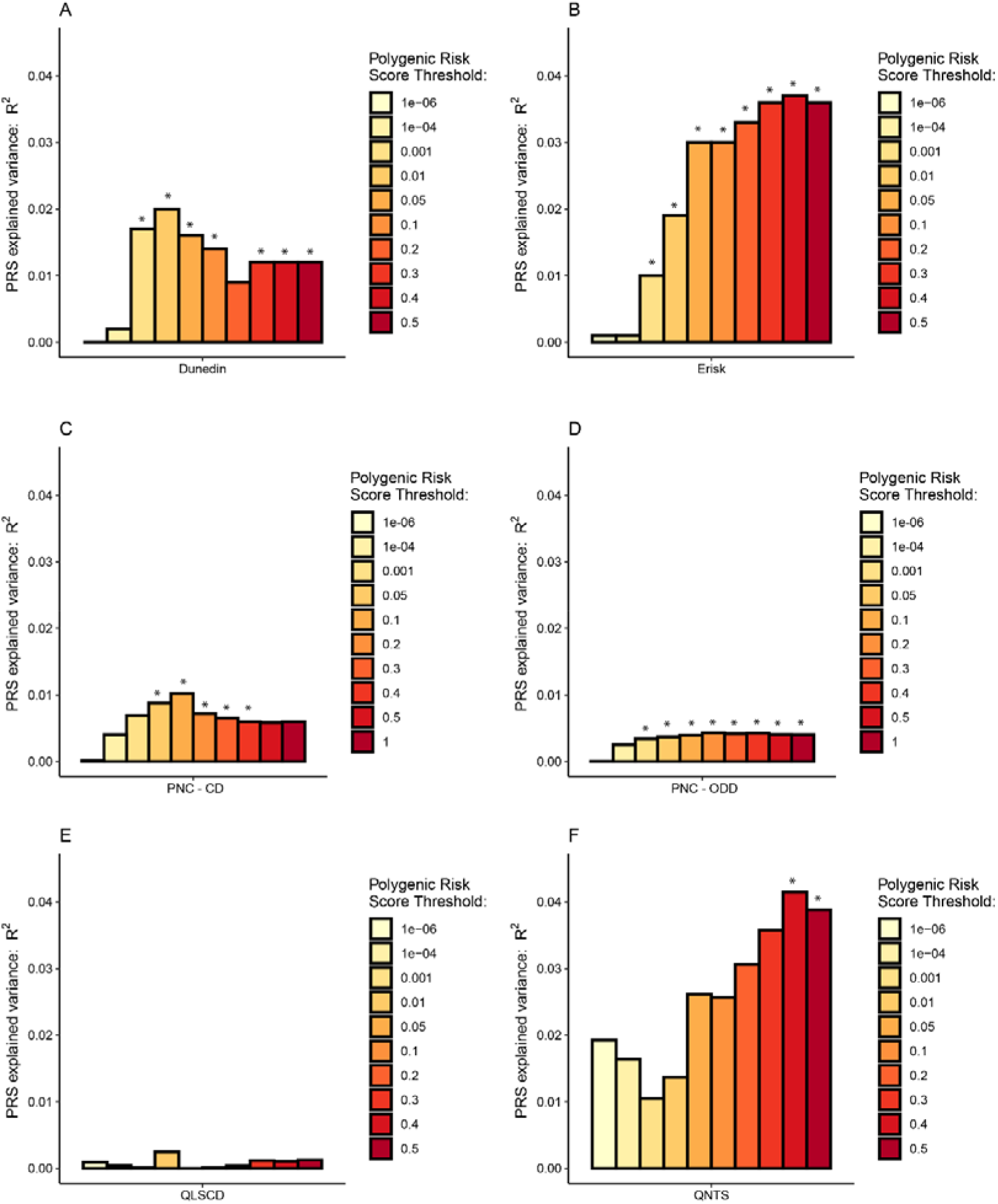
Polygenic risk score (PRS) associations of broad ASB with six antisocial outcomes in five cohorts. Bar charts illustrating the proportion of variance (incremental R^2^, or ΔR^2^) explained by the PRSs. PRSs are shown for broad ASB associated with childhood ASB in the Dunedin Longitudinal Study [A], with externalizing behavior in the E-Risk Study [B], with Conduct Disorder [C] and Oppositional Defiant Disorder [D] in the Philadelphia Neurodevelopmental Cohort Study, with ASB in the Quebec Longitudinal Study of Children’s Development Study [E], and with time-aggregated ASB in the Quebec Newborn Twin Study [F]. Asterisks (*) show statistical significance after applying a Bonferroni correction on the 22 tested phenotypes at P□<□0.0023.

**Figure 3:**
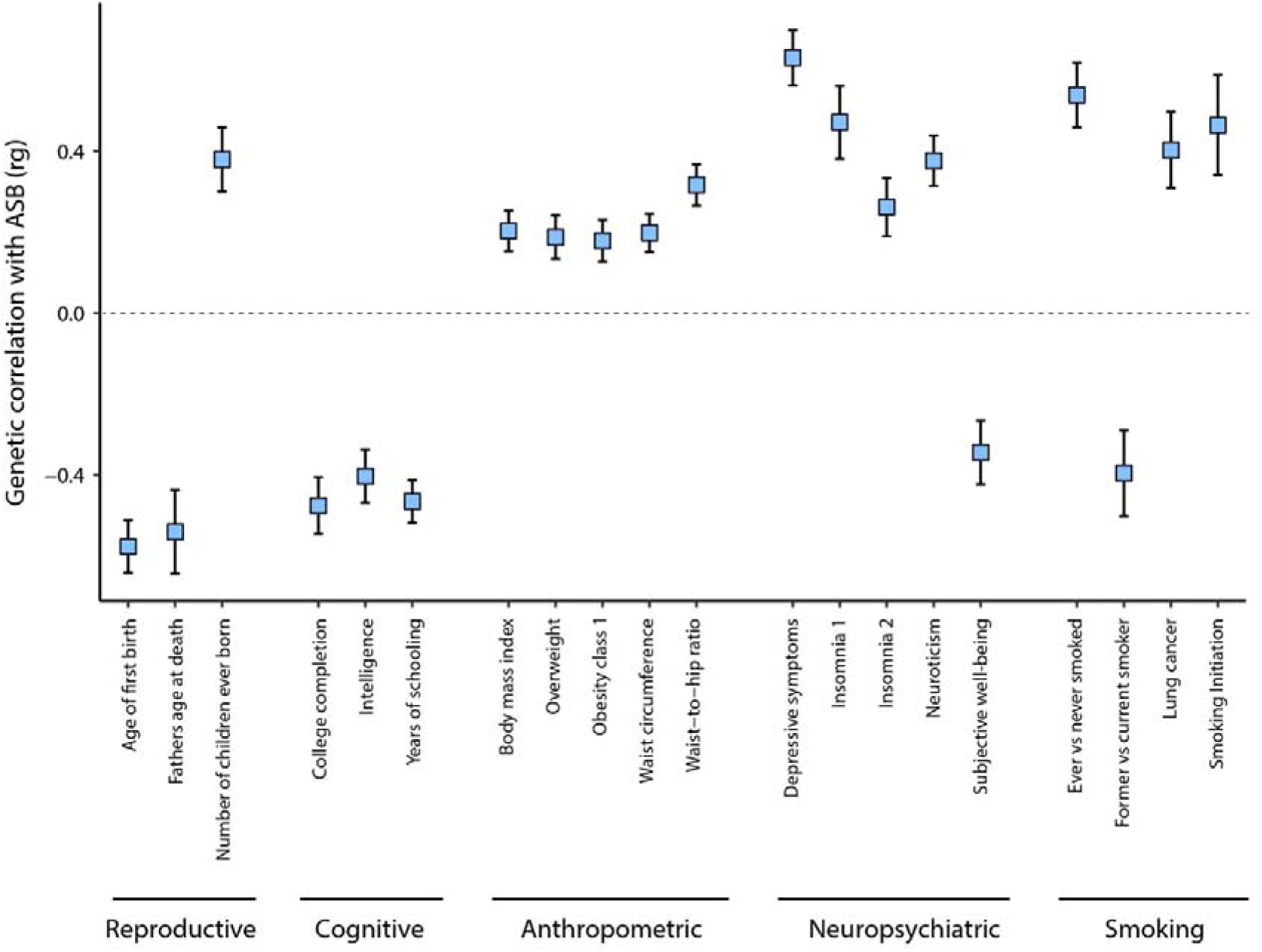
Genetic correlations of traits and diseases that were significantly associated with ASB. Significant genetic correlations of ASB with previously published results of other traits and diseases, computed using cross-trait LD Score Regression in LDHub, Bonferroni-corrected P-value: 0.00074 (bars represent 95% confidence intervals).

### Mouse model of pathological aggression

Whole genome sequencing analysis of SNVs in aggressive antisocial BALB/cJ mice compared to BALB/cByJ mice controls revealed differences between these lines located in introns of *Foxp2* (rs241912422) and *Cntnap2* (rs212805467; rs50446478; rs260305923; rs242237534), a well-studied neural target of this transcription factor.

### Heritability and Polygenic Scoring

#### SNP heritability

To assess the proportion of variance in liability for broad ASB explained by all measured SNPs, we computed the SNP-based heritability (h^2^_SNP_), which was estimated to be 8.3% (s.e. = 1.2%) by LD score regression (LDSC)^24^.

## Polygenic Risk Scoring in five independent cohorts

To assess how well the PRS derived from our ASB GWAS meta-analysis predicts other measures of antisocial behavior, we carried out PRS analyses in five independent cohorts (Supplementary Note 7).

### Dunedin Longitudinal Study

In New Zealand, participants were derived from the Dunedin Longitudinal Study^25^ (N=1,037, assessed 14 times from birth to age 45 years). We tested nine phenotypes and found significant associations with the BroadABC-based PRS for two: childhood ASB and official-records of juvenile convictions. Although not surviving Bonferroni adjustment, we found nominal significant (P < 0.05) association with the BroadABC-based PRS for eight phenotypes. We did not find evidence for a PRS association with partner violence. Lastly, we compared individuals grouped into the following four distinct developmental trajectories of antisocial behavior using general growth mixture modeling: low antisocial behavior across childhood through adulthood, childhood-limited antisocial behavior, adolescent-onset antisocial behavior, and life-course persistent antisocial behavior^26^. Individuals following the life-course persistent (LCP) antisocial trajectory were characterized by the highest levels of genetic risk (see Supplementary Figure 2); the nominally significant higher PRS of the LCP trajectory group compared to the low ASB group (P = 0.032 and P = 0.049, for P-value thresholds 0.05 and 0.1 respectively) did not survive Bonferroni adjustment. For a full report of the findings in the Dunedin cohort, see Supplementary Table 9 and Supplementary Note 8.

### Environmental Risk Longitudinal Twin Study (E-Risk)

In England and Wales, participants were included from the E-Risk Study (N=2,232, assessed five times from birth to age 18 years). We tested eight phenotypes and found significant associations for seven. PRS analyses revealed significant associations with parent- and teacher-reported antisocial behavior up to age 12 years, conduct disorder diagnosis up to age 12 years, with the externalizing spectrum at age 18 years, and with official records of criminal convictions up to age 22 years. For a full report of the findings in the E-risk Study, see Supplementary Table 10 and Supplementary Note 8.

### Philadelphia Neurodevelopmental Cohort (PNC)

In the United States, participants were included from the PNC Study (N=4,201). We tested two phenotypes and found significant associations for both. We found that higher PRS for ASB were associated with symptom counts of both conduct disorder (P < 0.0001, delta R^2^=1.0%, Supplementary Table 11) and oppositional defiant disorder (P < 0.0001, delta R^2^=0.4%, Supplementary Table 12).

### Quebec Longitudinal Study of Children’s Development (QLSCD)

In Canada, participants were included from the QLSCD study (N=599). We tested one phenotype and did not find a significant association (P > 0.05, Supplementary Table 13) between PRS and the score on a self-report questionnaire related to conduct disorder, delinquency, and broad antisocial behavior in young adults (age range= 18-19 years).

### Quebec Newborn Twin Study (QNTS)

In Canada, participants were derived from the QNTS study (N=341). We tested two phenotypes and found a significant association for one. We computed a factor score based upon five teacher-rated assessments of ASB in youngsters during primary school (age range= 6-12 years). We found that higher PRS were associated with a higher factor score of ASB (P = 0.001, for P-value thresholds .4, adjusted delta R^2^=3.9%, Supplementary Table 14). We failed to find evidence for an association between PRS and self-reported antisocial behavior in young adults (P > 0.05).

## Genetic correlations through LD score regression

ASB is known to correlate with an array of problems^17^. To test whether these phenotypic associations are also reflected in genetic correlations we performed analyses with LDSC in 68 traits and diagnoses (Supplementary Table 15). We found strong correlations between ASB and reproductive traits (e.g. younger age of first birth (rg□=□-0.58, s.e.□=□0.06, P□=□2.93□×□10^-15^)), cognitive traits (e.g. fewer years of schooling (rg□=□-0.49, s.e.□=□0.06, P□=□1.94□×□10^-10^)), anthropometric traits (e.g. increased waist-to-hip ratio (rg = 0.32, s.e. = 0.05, P□ = □5.59□×□ 10^-6^)), neuropsychiatric traits (e.g. more depressive symptoms (rg□ = □0.63, s.e.□=□0.07, P□ = □2.45□×□10^-16^)) and smoking related traits (e.g. ever smoked (rg□=□0.54, s.e.□=□0.08, P□=□1.48□×□10^-7^)).

## Discussion

Our GWAS meta-analysis of broad ASB in 85,359 individuals from population cohorts and those with a clinical diagnosis related to ASB, revealed one novel associated locus on chromosome 7 (7:114043159, rs12536335), residing in the forkhead box P2 *(FOXP2)* gene. The lead SNP is relatively proximal (~14kb upstream) to an important enhancer region located 330 kb downstream of the first transcriptional start site (TSS1) of the gene^20^. This SNP is also in the vicinity (~8kb upstream) of a second transcriptional start site (TSS2) of *FOXP2* that can drive expression of alternative transcripts. The *FOXP2* gene is expressed in sensory, limbic, and motor circuits of the brain, as well as the lungs, heart, and gut^20^. It encodes a transcription factor that acts as a regulator of numerous target genes and has been implicated in multiple aspects of brain development (e.g. neuronal growth, synaptic plasticity)^27^. *FOXP2* was first identified two decades ago when rare heterozygous mutations of the gene were linked to a monogenic disorder involving speech motor deficits, accompanied by impairments in expressive and receptive language^28,29^. Nevertheless, so far there is little evidence for a role of common *FOXP2* variants in interindividual differences in language function^30,31^. Thus, even though earlier behavioral research^32,33,34^ has reported a link between language problems and ASB, we should not over-interpret the *FOXP2* findings of the present study. Moreover, SNPs at this locus have been associated, through GWAS, with a range of externalizing traits, including ADHD^35^, cannabis use disorder^36^, and generalized risk tolerance^37^. Given the involvement of SNPS at this locus in different behavioral traits and diagnoses, and considering the small effect sizes, it is clear that the association of FOXP2 variation with ASB has limited explanatory value on its own, but could yield insights once placed in broader context by future research.

In the present study we also compared the BALB/cJ strain, a mouse model of pathological aggression, to BALB/cByJ controls, and found intronic variants in *Foxp2* and one of its downstream targets, *Cntnap2.* Previous studies in human cellular models have shown that the protein encoded by *FOXP2* can directly bind to regulatory regions in the *CNTNAP2* locus to repress its expression^38^. Interestingly, mice with cortical-specific knockout of *Foxp2* have been reported to show abnormalities in social behaviors^39^.Although these findings may indicate that the intronic SNVs are relevant to the behavioral differences between the strains, further evidence is needed to show that the variants actually have functional relevance for the mouse phenotype. Future studies may utilize complementary data comparing gene expression in the two mouse lines or could investigate functional impact (e.g. do they map to credible enhancer regions, are they likely to alter binding for transcription factors?) of the SNVs identified.

In contrast with the previous BroadABC GWAS analyses, we did not find evidence for sex-specific genetic effects in the present study. Although we did have access to sex-specific data in considerable subsets (N = 22,322 males, N = 26,895 females), the power to detect new variants employing such sample sizes is still limited. Compared to our previous study, we found that the variance explained in independent samples by PRS based on the resulting summary statistics has substantially increased from 0.21% to 3.9%. Essentially, we found consistent links of our ASB PRS with multiple antisocial phenotypes at different developmental stages, from different reporting sources, and reflecting measurements from different disciplines (psychology, psychiatry, criminology). These links were found in individuals from New Zealand, Britain, the United States, and Canada, born 30 years apart. We also show that our ASB PRS were more strongly associated with more severe and persistent types of ASB.

Notwithstanding the increase of effect size of the PRS, and calculations yielding a more precise estimate, the variance explained by the PRS was still relatively small, which was expected in light of the low SNP heritability of 8.3%. Given the highly polygenic architecture of ASB, contributing SNPs have low average effect sizes, thus leading to limited predictive power in independent samples. New PRS methods along with further increasing sample sizes will likely further increase the amount of variance accounted for by the PRS. Moreover, the association may be enhanced by improving the quality of phenotype measurements, which is reflected by our PRS results demonstrating the most robust association with high quality measurement of ASB (using a factor score based upon multiple assessments). Aggregating data from measurements across ages, as opposed to the measures assessed at a single time point, can lead to more reliable trait measures and to better prediction^40^. Phenotypically, adding more extreme ASB phenotypes to the GWAS meta-analysis might also lead to more explained variance. Thus, future efforts of the BroadABC will continue to focus on more severe forms of ASB and its persistence across the lifespan. Moreover, by considering genetically correlated traits through multi-trait GWAS methods^41^ and multi-trait PRS methods^42^ it might be possible to boost power for discovery through GWAS meta-analysis and PRS prediction. Lastly, a major limitation of the present study is that our GWAS results are limited to individuals of European ancestry. This Eurocentric bias may lead to more accurate predictions in individuals with European ancestry, compared to non-Europeans, thus potentially increasing disparities in outcomes related to ASB^43,44^. To realize the full and equitable potential of polygenic risk, future genetic studies on ASB should also include non-European samples.

Developmental criminological research findings, such as the influential developmental taxonomy theory by Moffitt^45,46^, have established the existence of distinctive offending patterns across the life-course^47^. These distinctive developmental trajectories of ASB are thought to have different underlying etiological processes, with higher genetic influences for life-course-persistent offending as compared to the more socially influenced adolescence-limited offending. Barnes and coworkers showed that the heritability was not uniform across different offending groups, suggesting that the causal processes may vary across offending patterns^48,49^. In the present study we found a trend of higher PRS for ASB showing a stronger association with the life-course-persistent trajectory of ASB as compared to the low ASB group. The life-course-persistent trajectory is also known to be associated with the most profound brain alterations and poorest brain health^50^. These findings are important since they can improve understanding of downstream neurobiological mechanisms relevant to the etiology of antisocial development^50^. Sufficiently powered future studies should thus aim to further elucidate the genetic risk and protective factors that underlie different offending trajectories^51^.

Our genetic correlation analyses confirmed previously reported^13,17,52^ correlations between ASB and a wide range of traits and diagnoses. Partial sharing of genetic effects does not necessarily represent causal relationships, yet merely signifies the presence of potentially shared biology or other mechanisms linking the conditions^53^. Therefore, it is likely that there are common underlying genetic factors increasing general vulnerability to psychopathologies. These comorbid effects are in line with findings in the Dunedin Study demonstrating that life-course-persistent offenders are characterized by several pathological risk factors, related to domains of parenting, neurocognitive development, and temperament^46^. This signifies the importance of investigating pleiotropy and considering the complex etiology of the broader ASB phenotype. Large-scale collaborations, such as the BroadABC, will facilitate the expansion of epidemiological studies capable of further exploring the interaction of genetic risk and socio-environmental risks, and how these contribute to the multifaceted origin of ASB.

## Methods

### Samples

The meta-analysis included 21 new discovery samples of the BroadABC with GWAS data on a continuous measure of ASB, totaling 50,252 participants: The National Longitudinal Study of Adolescent to Adult Health^54^ (ADH), Avon Longitudinal Study of Parents and Children^55–57^ (ALSPAC), Brain Imaging Genetics^58^ (BIG), CoLaus-PsyCoLaus^59^, Collaborative Study on the Genetics of Alcoholism^60^ (COGA), Finnish Twin Cohort^61^ (FinnTwin), The Genetics of Sexuality and Aggression^62^ (GSA), Minnesota Center for Twin and Family Research^63^ (MCTFR), Phenomics and Genomics Sample^64^ (PAGES), eight samples of the QIMR Berghofer Medical Research Institute (QIMR; 16Up project [16UP^65^], Twenty-Five and Up Study [25UP^66^], Genetics of Human Agency [GHA^67^], Prospective Imaging Study of Ageing [PISA^68^], Semi-Structured Assessment for the Genetics of Alcoholism SSAGA Phase 2 [SS2^69^], Genetic Epidemiology of Pathological Gambling [GA^70^], Twin 89 Study [T89^71^], and Nicotine Study [NC^72^]), Spit for Science^73^ (S4S), two samples (from different genotype platforms) of the Twin Early Development Study^74^ (TEDS), and the TRacking Adolescents’ Individual Lives Survey^75^ (TRAILS).

We complemented the above data with GWAS summary statistics on case-control data on disruptive behavior disorders from the recently published Psychiatric Genetics Consortium/iPSYCH consortium meta-analysis, which included data from seven cohorts (Cardiff sample, CHOP cohort, IMAGE-I & IMAGE-II samples, Barcelona sample, Yale-Penn cohort, and the Danish iPSYCH cohort), totaling 3,802 cases and 31,305 controls^16^.

We observed a high genetic correlation between the 21 meta-analyzed BroadABC samples and the 7 Psychiatric Genetics Consortium/iPSYCH samples, with the ‘Effective N’ as weight (*r*_g_□ = ≥ 0.93, P = 9.04 ×□ 10^-8^), indicating strong overlap of genetic effects. Hence, we continued with the combined 28 samples (N= 85,359) for all analyses.

All included studies were approved by local ethics committees, and informed consent was obtained from all of the participants. All study participants were of European ancestry. Full details on demographics, measurements, sample analysis, and quality control are provided in Supplementary Table 1.

### Genome-wide association analysis and quality control of individual cohorts

In all 28 discovery samples, genetic variants were imputed using the reference panel of the Haplotype Reference Consortium (HRC) or the 1000G Phase 1 version 3 reference panel. The regression analyses were adjusted for age at measurement, sex, and the first ten principal components. To harmonize the imputation, data preparation, and genome-wide association (GWA) analyses, a specific analysis protocol (Supplementary Note 1) was followed in the 18 BroadABC discovery samples. Further details on the genotyping (platform and quality control criteria), imputation, and GWA analyses for each cohort are provided in Supplementary Table 2.

Two semi-independent analysts (JJT & EU) performed stringent within-cohort quality control, filtering out poor performing SNPs. SNPs were excluded if they met any of the following criteria: study-specific minor allele frequency (MAF) corresponding to a minor allele count (MAC)□<100, poor imputation quality ((INFO/R2) score□<0.6), and/or Hardy-Weinberg equilibrium P□<□5□×□10^-6^. Moreover, we excluded SNPs and indels that were ambiguous (A/T or C/G with MAF□>0.4), duplicated, monomorphic, multiallelic, or reference-mismatched (Supplementary Note 2, Supplementary Table 17). Then, we visually inspected the distribution of the summary statistics by creating quantile-quantile plots and Manhattan plots for the cleaned summary statistics from each cohort (Supplementary Notes 4, 5 and 6). Discrepancies between the results files of the two semi-independent analysts were examined and errors corrected.

### Meta-analyses on combined and sex-specific samples

A meta-analysis of the GWAS results of the 28 discovery samples (N = 85,359) was performed through fixed-effects meta-analysis in METAL, using SNP P-values weighted by sample size. After combining all cleaned GWAS data files, meta-analysis results were filtered to exclude any variants with N□<□30,000. Consequently, we removed 2,134,049 SNPs, resulting in 7,392,849 SNPs available for analysis. To investigate sex-specific genetic effects, we also ran the meta-analysis in the datasets for which we had sex-specific data (N = 50,252). However, sex-specific SNP heritabilities, as estimated with LD Score Regression, were small and non-significant (3.7% (s.e. = 2.2%) for males and 1.0% (s.e. = 1.8%) for females). Due to the non-significant sex-specific heritability estimates, the genetic correlation of male and female ASB could not be estimated reliably and no sex-specific follow-up analyses were conducted.

### Whole-genome sequencing based on genetic differences between the BALB/c strains

Through whole-genome sequencing, we identified single nucleotide variants that distinguish aggressive BALB/cJ mice from control BALB/cByJ strains^76^. Sequencing libraries were prepared from high-quality genomic DNA using the TruSeq DNA PCR-Free kit (Illumina) and ultra-deep whole genome sequencing (average 30X read-depth across the genome) was performed on a HiSeq X Ten System (Illumina). We developed an efficient data processing and quality control pipeline. Briefly, raw sequencing data underwent stringent quality control and was aligned to either the mm10 (BALB/cJ versus BALB/cByJ strain comparison). Isaac^77^ was used to align reads and call single nucleotide variations (SNVs). We excluded SNVs that were covered by less than 20 reads, and that were not present in both animals from the same strain. SnpEff^78^ was used to annotate SNVs and explore functional effects on gene function. SNVs differing between the two strains were annotated to a total of 1573 genes, which were subdivided into three different categories (intronic/exonic noncoding and synonymous variants (1422 genes), untranslated regions (90 genes), missense mutations and splicing variants (61 genes)).

### Polygenic Risk Score Analyses

Polygenic risk scores (PRS) were created for ASB using all available SNPs of the discovery dataset^79,80^. PRS were computed as the weighted sum of the effect-coded alleles per individual. We calculated the PRS for subjects of five independent datasets, selected for their detailed phenotypes related to antisocial outcomes: (1) the Dunedin Study^40^, (2) the E-risk study^81^, (3) the Philadelphia Neurodevelopmental Cohort^82^, (4) the Quebec Longitudinal Study of Child Development^83^, and (5) the Quebec Newborn Twin Study^84^. All individuals were of European ancestry. To maintain uniformity across target cohorts, we adhered to the following parameters: Clumping was performed by removing markers in linkage disequilibrium, utilizing the following thresholds: maximum r2 = 0.2, window size = 500 kb. We excluded variants within regions of long-range LD^85^ (including the Major Histocompatibility Complex, see Supplementary Table 16 for exact regions). Second generation PLINK^86^ was employed to construct PRS for each phenotype, at the following 10 thresholds: P□<□1□×□10^-6^, P□<□1□×□10^-4^, P□<□1□×□10^-3^, P□<□1□×□10^-2^, P□ < 0□.05, P□<□0.1, P□<□0.2, P□<□0.3, P□< 0□.4, PL<10.5. To correct for multiple testing, we applied a Bonferroni correction on the 22 tested phenotypes (α = 0.00227).

### Genetic correlation analysis

To estimate the genetic correlation between ASB and a range of other phenotypes, we employed Linkage Disequilibrium Score Regression (LDSC)^24^ through the LD Hub web portal (http://ldsc.broadinstitute.org/ldhub/)^87^. We corrected for multiple testing by applying a Bonferroni correction on the 68 tested genetic correlations (α = 0.0007).

## Acknowledgements

G.M-V. was supported by a Doctoral Research Scholarship from the FRQSC. M.B., I.O-M., and J-P Gouin are supported by the Canada Research Chair Program. K.R. is supported by a Sir Henry Wellcome Postdoctoral Fellowship. ADB and DD were supported by grants from the Lundbeck Foundation (R102-A9118, R155-2014-1724 and R248-2017-2003), the EU FP7 Program (Grant No. 602805, “Aggressotype”) and H2020 Program (Grant No. 667302, “CoCA”), NIMH (1U01MH109514-01. JL, AM, and KR acknowledge the following funders: Academy of Finland, the Signe and Ane Gyllenberg foundation, Juho Vainio foundation, Yrjö Jahnsson foundation, Jalmari and Rauha Ahokas foundation, Sigrid Juselius Foundation and The Finnish Society of Sciences and Letters. DD is supported by NIH R01 AA015416 (Finnish Twin Study), P50 AA022537 (Alcohol Research Center), R25 AA027402 (VCU GREAT), R34 AA027347 (Personalized Risk Assessment), R01 AA028064 (Parental Marital Discord, PI: JS), and U10 AA008401 (COGA) from the National Institute on Alcohol Abuse and Alcoholism (NIAAA), and by R01 DA050721 (Externalizing Consortium) from the National Institute on Drug Abuse (NIDA). BF, MK, and NRM were also supported by funding from the European Community’s Horizon 2020 Programme (H2020/2014 – 2020) under grant agreements n° 728018 (Eat2beNICE) and n° 847879 (PRIME). They also received relevant funding from the Netherlands Organization for Scientific Research (NWO) for the Dutch National Science Agenda NeurolabNL project (grant 400-17-602). HMS and MRM are members of the Medical Research Council (MRC) Integrative Epidemiology Unit at the University of Bristol (MC_UU_00011/7). This work is also supported by the NIHR Biomedical Research Centre at University Hospitals Bristol NHS Foundation Trust and the University of Bristol. The views expressed in this publication are those of the authors and not necessarily those of the NHS, the National Institute for Health Research or the Department of Health and Social Care. AW has been supported by funding from the European Union Seventh Framework Programme (FP7/2007-2013) under grant agreement no. 602768 for the ACTION consortium. JK has been supported by the Academy of Finland Academy professorship program (grants 265240 and 263278). LC-C was supported by a QIMR Berghofer Research Fellowship. JJM was supported by a QIMR Berghofer International PhD Scholarship. ML was funded by an NHMRC Boosting Dementia Leadership Fellowship (APP1140441). SEM and NGM were funded by NHMRC investigator grants (APP1172917 and APP1172990).

Additional acknowledgements for each study cohort are described in **Supplementary Table 1**.

