## Supplementary Information for "Uncovering the Genetic Architecture of Broad Antisocial Behavior through a Genome-Wide Association Study Meta-analysis"

### 1. Imputation and GWAS protocol for meta-analysis on antisocial phenotypes

This appendix lists the instructions that were send to all participating cohorts to harmonize the imputation, data preparation and GWA analyses on antisocial phenotypes.

Standardization of the procedures is very important, as it will increase the precision of the meta-analyses across all samples of the BroadABC.

**Content of this document:**

1. Instructions for phenotype and covariate coding
2. Instructions for genotype imputation
3. Analysis outline
4. Instructions on the format of the input files for the Meta-analysis

#### 1. Instructions for phenotypes and covariate coding

*Inclusion*
We propose to limit the analyses to subjects from European ancestry. Please let us know if you have a large group of individuals of non-European ancestry, as we may meta-analyze non-European samples as an additional project.

*Antisocial phenotypes*

Please perform the analyses on continuous data. If you have multiple measures at different ages with the same measurement instrument, please contact Jorim Tielbeek at to discuss which instrument(s)/age group(s) is most relevant for the consortium.

*Covariates - Please use these variables as a covariate in your GWAS analysis:*

- **Age** at the time of the phenotypic assessment (in years since birth).
- **Sex** coded as 1=male, 0=female.
- **Population structure**. Please use the first ten principal components to correct for population structure in your sample. If necessary add study-specific covariates such as study site or batch effects.

#### 2. Instructions for genotype imputation

* If your data is already imputed to HRCv1.1, you can ignore this step and proceed to section 3 of the protocol.

Imputation through an imputation server is useful since it performs automatic quality checks (SNP names, alleles, MAFs etc.) and makes certain that imputations across cohorts are done to the exact same reference panel.

We recommend performing imputations using the reference panel of the Haplotype Reference Consortium (HRC). The HRC has assembled 64,976 haplotypes by combining sequence data from multiple cohorts of predominantly European ancestry. Imputation through the Sanger or Michigan imputation servers works relatively straightforward and results in a large number of testable genetic variants, thereby yielding an increased power for GWAS (compared to 1000Genomes).

If it is not possible to impute genotypes to HRC due to technical or privacy constraints, then summary statistics of analyses performed on genotypes imputed to 1000Genomes (phasev3 or higher) are also sufficient.

*Preprocess your genotype data before imputation*

Make sure your genotype data is mapped to NCBI Build 37 before starting the imputation.

We have specified below the typical pre-imputation QC steps that we recommend. Analysts may of course decide to perform additional QC steps. QC can be performed in Plink. Please provide a brief description of QC criteria performed in your cohort.

- Relatedness (kinship coefficient > 0.05), unless part of a twin/family study design where relatedness is controlled for in the genetic analysis
- Sample call rate (cut off >95% threshold recommended)
- Exclude samples with heterozygosity > median + 3*IQR
- Remove gender mismatches
- Remove duplicates
- Remove PCA outliers using a PCA projection of the study samples onto 1KG reference samples.
- Hardy-Weinberg p>10-6, SNP call rate ≥98%
- Remove monomorphic markers
- Remove ambiguous (A/T and C/G) SNPs

Additional QC will be conducted centrally at the meta-analysis stage.

*Choose an imputation server*

There are currently two imputation servers available to impute to the HRC reference panel:

- Sanger Imputation server (https://imputation.sanger.ac.uk/)
- Michigan Imputation server : (https://imputationserver.sph.umich.edu/index.html)

Note that standard encryption (SSL or SFTP) is used on both servers to ensure that genotypes are encrypted during upload and download.

*Upload your VCF files on the server*

Create an account, login and follow the instructions on the website. Note that we use the Michigan server as an example, the Sanger server works similarly.

Since only VCF files can be submitted, you will need to convert your files to this format. If your genotype files are currently in binary plink format (.bed, .bim, .fam) you can convert them using the following command (from current working directory):

plink --bfile <binary plink prefix> --recode vcf-iid --out vcf_input_file_prefix

Next, you should use bgzip to make a zipped archive of the newly created .vcf file, as the imputation server only accepts genotype data zipped by bgzip, command:

bgzip filename.vcf

Now you can upload the VCF files. Then select the following options:

- Reference Panel: HRC r1.1 2016
- Phasing: Eagle v2.3
- Population: EUR (this parameter is for quality control purposes)
- Mode: Quality Control & Imputation

*Downloading your imputed genotypes, info files, and QC report*

When imputation has finished you will be notified by email. The imputation server will automatically encrypt all your imputed genotypes (for protection during download). The password to decrypt the files will be in the email notification, so don’t delete that email!

Please download all available files (the QC report, statistics, zip files, and all the log files), since we may need them for future quality checks. We also ask you to quickly check the qcreport.html and statistics.txt files before proceeding with the analysis plan. Should you be unsure of your imputation quality, please do not hesitate to contact us.

Additional information regarding preparation of your data for imputation and running the imputation on the server is provided at the help page:
https://imputationserver.sph.umich.edu/index.html#!pages/help

#### 3. Analysis outline

In order to maximize power, continuous traits are preferred. If continuous data are not available, such as in some clinical cohorts with matched controls, please perform the analysis on the dichotomous trait.

Please perform the analyses for:

1. Only males
2. Only females
3. Combined

If you have two selected measurement instruments (e.g. SDQ and APSD), relevant for the consortium please run the three analyses for the two instruments separately (so six analyses in total).

*Continuous/quantitative:*

For continuous measures please use linear Regression onto estimated dose (as included in PLINK, MACHQTL, ProbABEL, SNPTEST, MERLIN), while adjusting for population structure and covariates.

Model Linear Regression: ASB score ~ sex + age + PCs + SNP

- SNP = genotype (estimated dose from 0 to 2)
    - Sex: coded 1 for male, 0 for female
    - Age: at measurement (in years)
    - PC's: principle components* (ancestry)

*Dichotomous/diagnostic:*

In case you only have case control status, please use logistic regression with dichotomous outcome. Please create a dichotomized score using the specific cutoff per measurement instrument, if no standard (such as DSM-IV/V) criteria exist, please contact Jorim Tielbeek to discuss.

Model Logistic Regression: ASB (dichotomous measure) ~sex + age + PCs + SNP

- SNP = genotype (estimated dose from 0 to 2)
    - Sex: coded 1 for male, 0 for female
    - Age: at measurement (in years)
    - PC's: principle components* (ancestry)

#### 2. Quality Control of discovery cohorts and meta-analysis

We meta-analyzed genome-wide association studies of ASB measures in 28 cohorts, which are described in detail in Supplementary Table 1 and 2.

In all 28 discovery cohorts, genetic variants were imputed using the reference panel of the Haplotype Reference Consortium (HRC) or with the 1000G Phase 1 version 3 reference panel. The regression analyses were adjusted for age at measurement, sex and the first ten principal components. To harmonize the imputation, data preparation and GWA analyses, a specific analysis protocol (Supplementary Note 1) was followed in the 18 BroadABC discovery samples. Further details on the genotyping (platform and quality control criteria), imputation and GWA analyses for each cohort are provided in Supplementary Table 2.

Two semi-independent analysts (JJT & EU) performed stringent within-cohort quality control, filtering out poor performing SNPs. SNPs were excluded if they met any of the following criteria: study-specific minor allele frequency (MAF) corresponding to a minor allele count (MAC) <100, poor imputation quality ((INFO/R2) score <0.6) or Hardy–Weinberg equilibrium $P < 5 \times 10^{-6}$. Moreover, we excluded SNPs and indels that were ambiguous (A/T or C/G with MAF >0.4), duplicated, monomorphic, multiallelic or reference mismatched. Then, we visually inspected the distribution of the summary statistics and created quantile–quantile plots and Manhattan plots for the cleaned summary statistics from each cohort (Supplementary Notes 4, 5 and 6). Discrepancies between the results files of the two semi-independent analysts were examined and errors corrected.

A meta-analysis of the GWAS results of the 24 discovery samples (N = 85 359) was performed through fixed-effects meta-analysis in METAL, using SNP P values weighted by sample size.

After combining all cleaned GWAS data files, meta-analysis results were filtered to exclude any variants with N < 30,000. Consequently, we removed 2,134,049 SNPs, resulting in 7,392,849 SNPs available for analysis. To investigate sex-specific genetic effects, we also ran the meta-analysis in the datasets for which we had sex-specific data (N = 50 252). However, sex-specific SNP heritabilities, as estimated with LD Score Regression, were small and non-significant (3.7% (SE = 2.2%) for males and 1.0% (SE = 1.8%) for females). Due to the non-significant sex-specific heritability estimates, the genetic correlation of male and female ASB could not be estimated reliably and no sex-specific follow-up analyses were conducted.

#### 3. Gene and Gene-Set Analyses

The statistical software tools MAGMA[1] and FUMA[2] were employed to perform the gene and gene-set analyses. We utilized the quality-controlled summary statistics of the combined meta-analysis results (N=85,359). The SNP p-values were used as input for MAGMA (which does not require access to the raw genotype data). We tested the accumulated association of multiple SNPs for 18,247 protein coding genes, by calculating gene-based test statistics as the mean of the chi-square statistics of all SNPs that reside within a gene, including a window of 50 kb around the gene. The genotype data of the 1,000 Genomes European reference panel were obtained to estimate the Linkage Disequilibrium (LD) between SNPs to account for LD within the genes[3]. Next, 10,678 gene sets (curated gene sets: 4761, Gene Ontology terms: 5917) from the Molecular Signatures Database (MSigDB[4], v6.2) were used to perform a gene-set analysis in MAGMA (which tests whether the genes in a gene set are more strongly associated with ASB than other genes in the genome), in which the gene analysis results served as input. The gene-set analysis corrects for gene size and gene density, and accounts for LD between genes using a gene correlation matrix based on the genotype data of the 1,000 Genomes European reference panel. Moreover, to test for association we with tissue-specific gene-sets we linked them to our gene-based p-values in FUMA.

Gene analyses in MAGMA identified the forkhead box protein P2 (*FOXP2*) gene to be associated with broad antisocial behavior ($P$ = 1.01 x $10^{-9}$, Supplementary Figure 1, Supplementary Table 6). Gene-set analyses did not reveal significant gene sets (Supplementary Table 7), and we also found no significant association results for the tissue-specific gene expression (Supplementary Table 8).

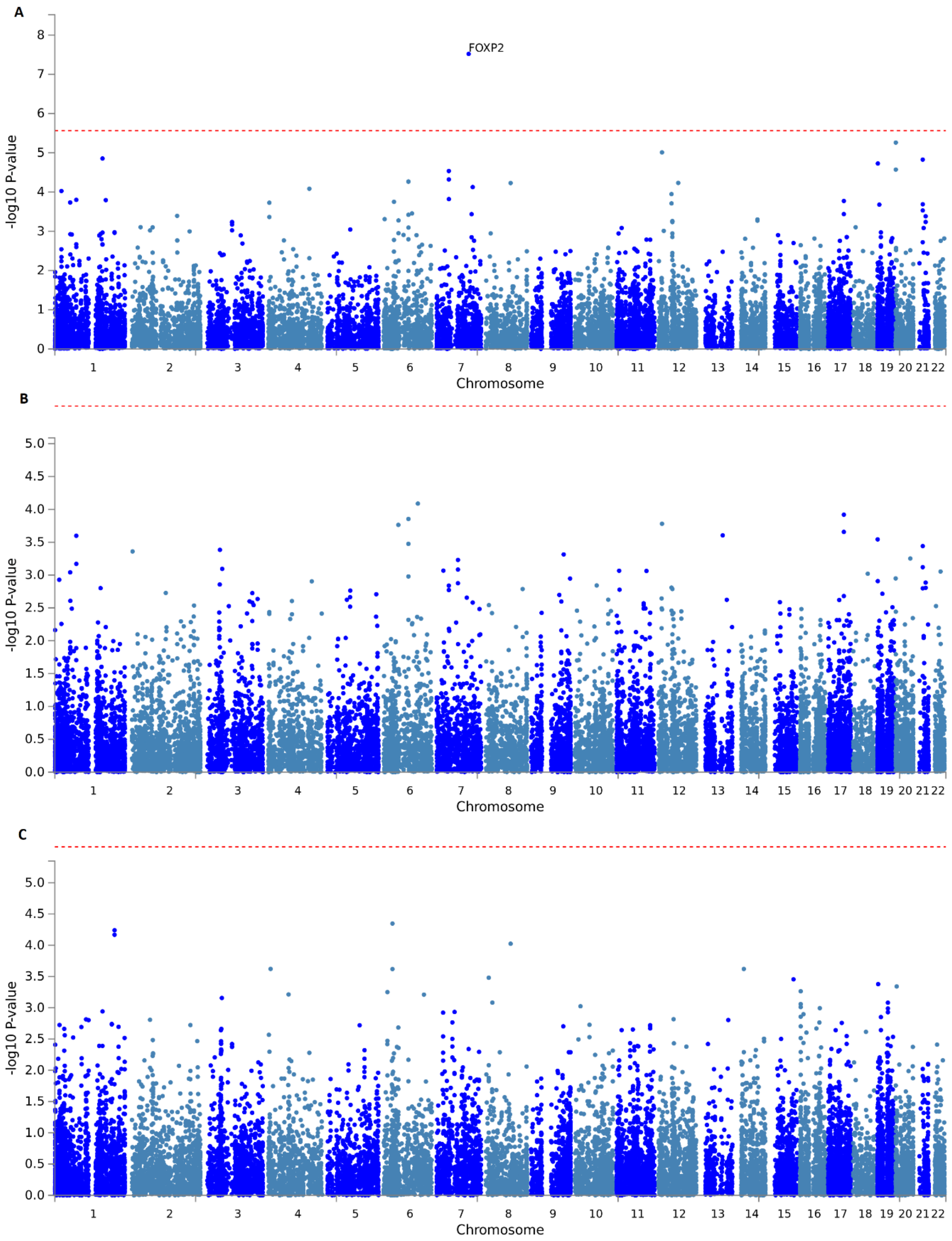

Supplementary Figure 1**.** Manhattan plot of the Genome Wide Gene Association Analysis for combined (upper panel, A), male-specific (middle panel, B) and female-specific (lower panel, C) meta-analyses. Input SNPs were mapped to 18,247 protein coding genes. Negative log10-transformed P-values for each gene are plotted. Gene-wide statistical significant associations (red dashed line in the plot) were defined at P = 0.05/18247 = 2.740e-6.

### 4. Manhattan and Q-Q plots discovery cohorts – only males

#### Manhattan plots

ADH

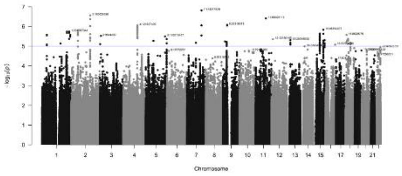

GSA

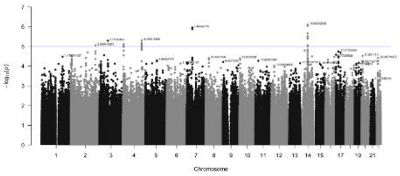

QIMRB_4

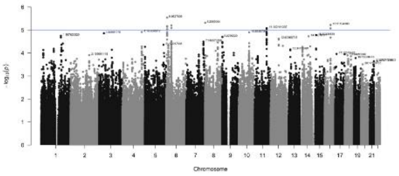

ALSPAC

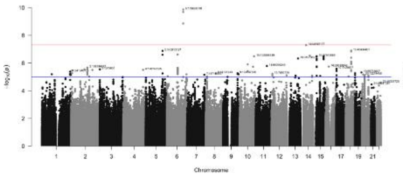

MCTFR

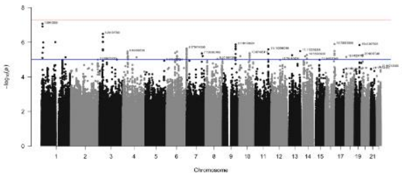

QIMRB_5

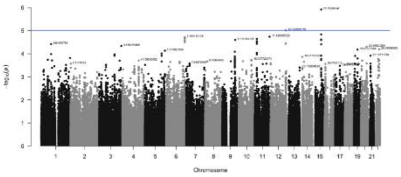

BIG

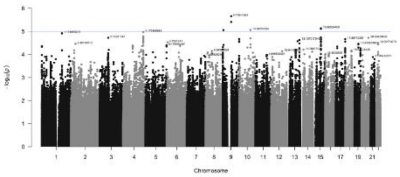

PAGES

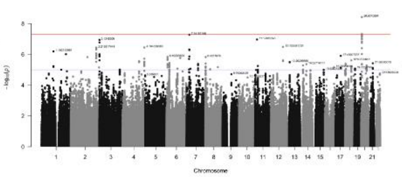

S42

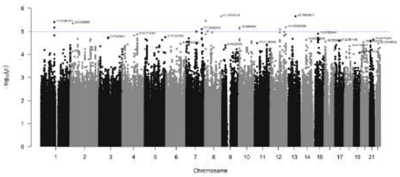

COGA

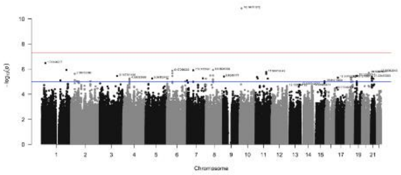

QIMRB_1

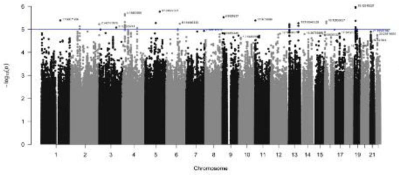

TEDS_1

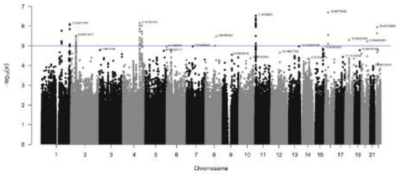

CoLaus

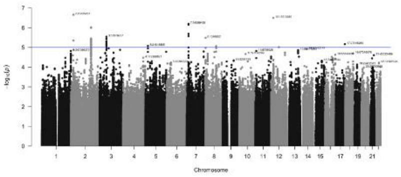

QIMRB_2

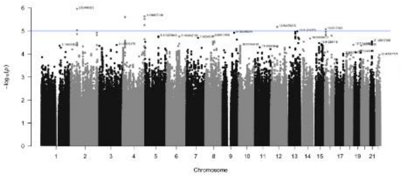

TEDS_2

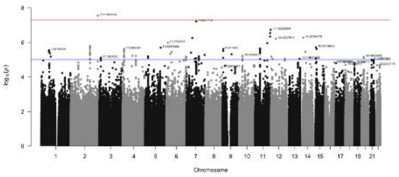

FinnTwin

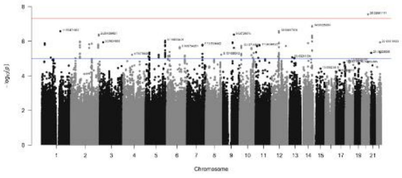

QIMRB_3

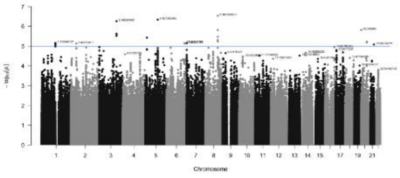

TRAILS

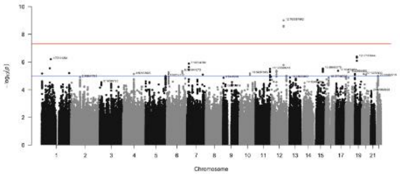

#### Quantile-quantile plots

ADH

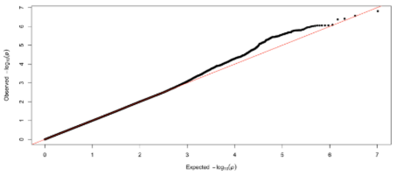

GSA

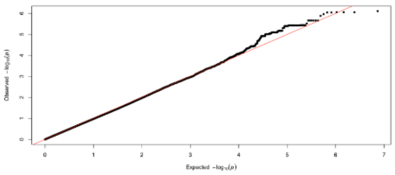

QIMRB_4

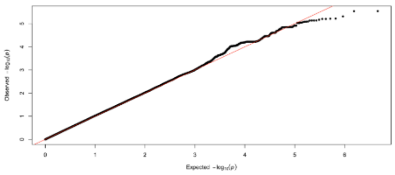

ALSPAC

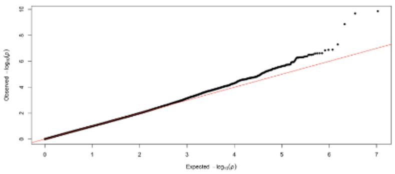

MCTFR

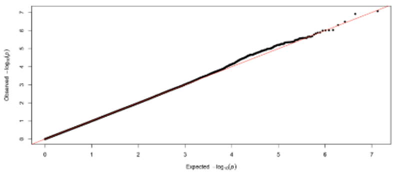

QIMRB_5

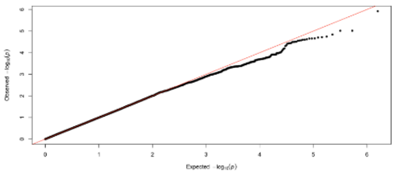

BIG

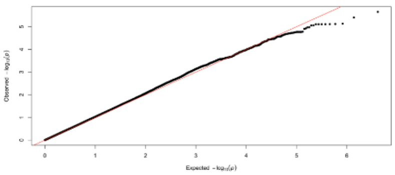

PAGES

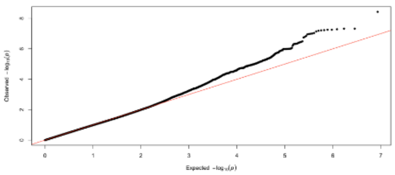

S42

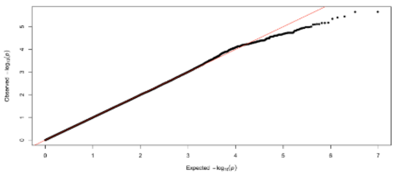

COGA

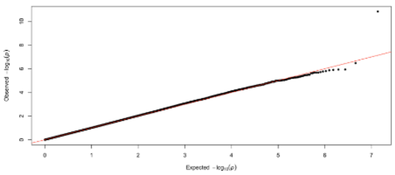

QIMRB_1

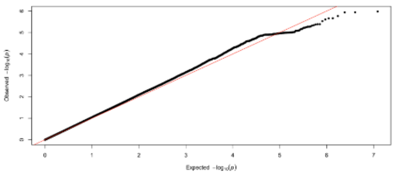

TEDS_1

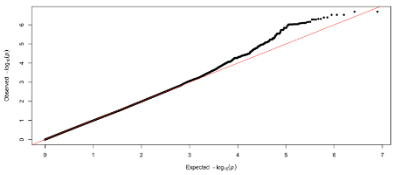

CoLaus

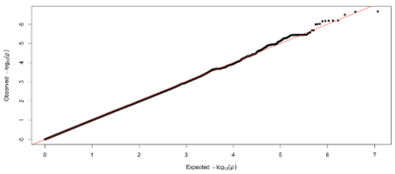

QIMRB_2

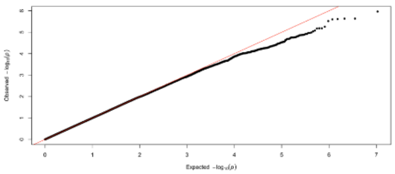

TEDS_2

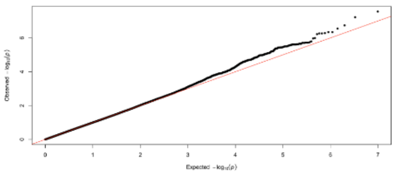

FinnTwin

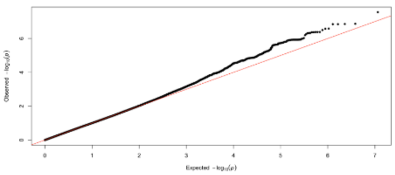

QIMRB_3

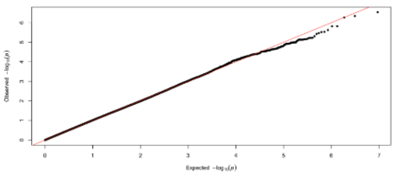

TRAILS

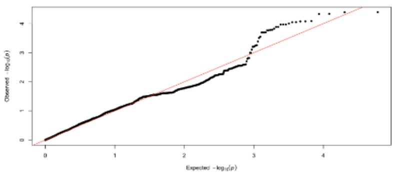

### 5. Manhattan and Q-Q plots discovery cohorts – only females

#### Manhattan plots

ADH

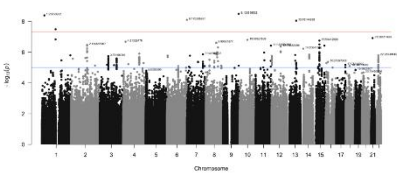

GSA

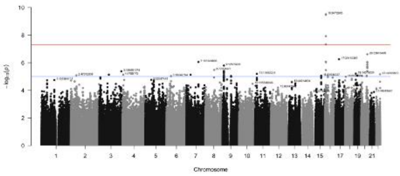

QIMRB_5

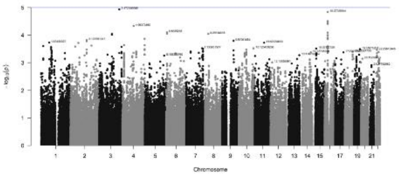

ALSPAC

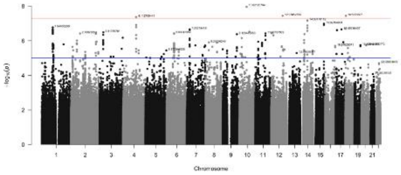

MCTFR

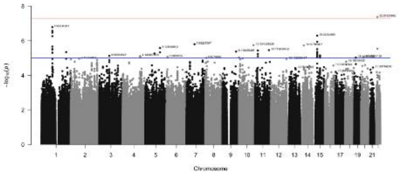

S42

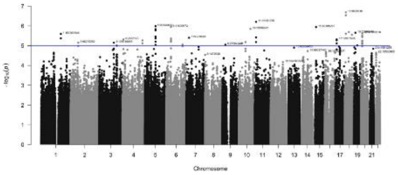

BIG

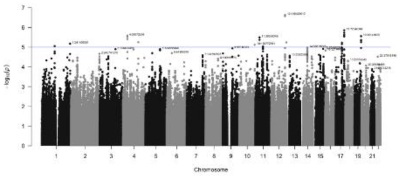

PAGES

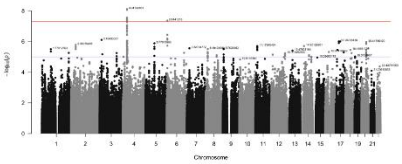

TEDS_1

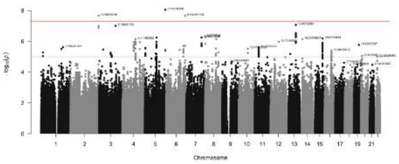

COGA

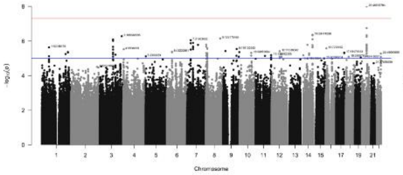

QIMRB_1

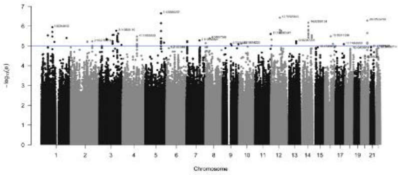

TEDS_2

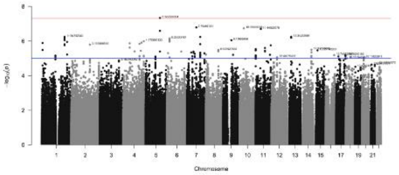

CoLaus

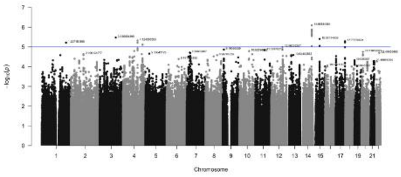

QIMRB_3

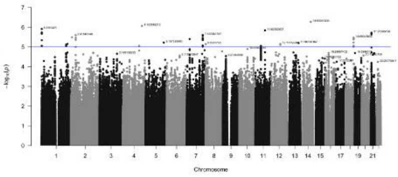

TRAILS

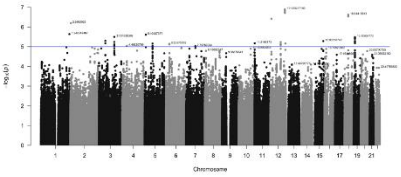

FinnTwin

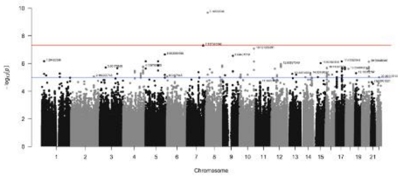

QIMRB_4

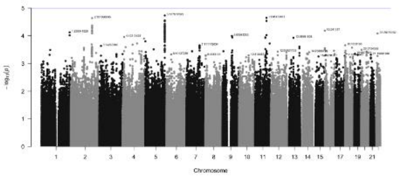

#### Quantile-quantile plots

ADH

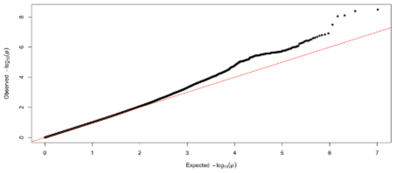

GSA

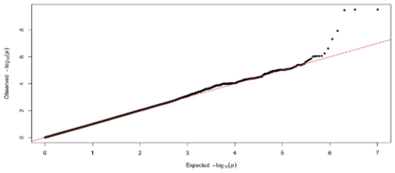

QIMRB_5

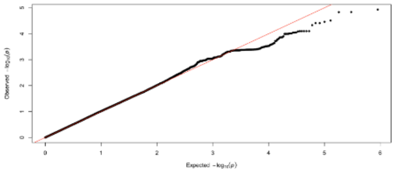

ALSPAC

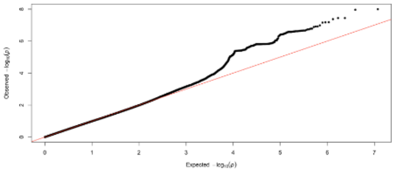

MCTFR

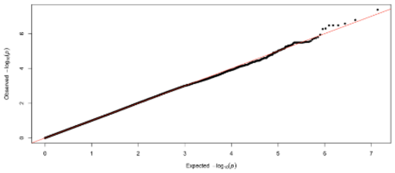

S42

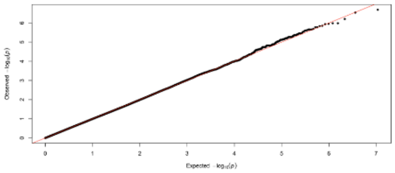

BIG

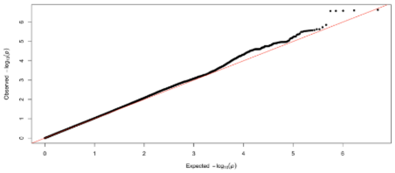

PAGES

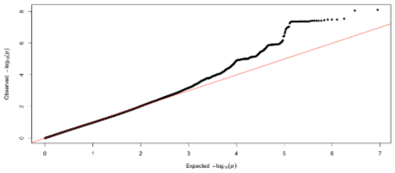

TEDS_1

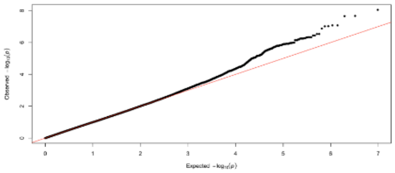

COGA

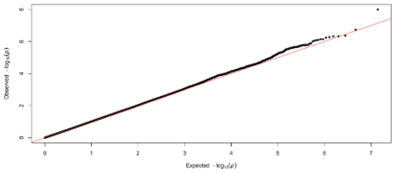

QIMRB_1

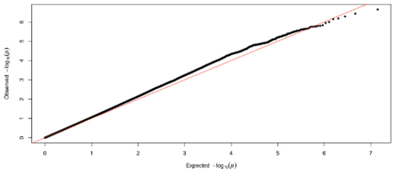

TEDS_2

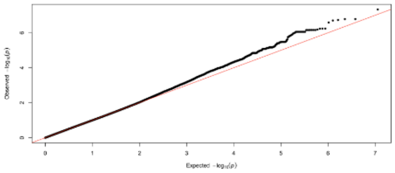

CoLaus

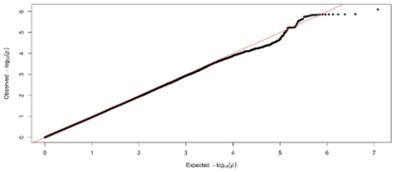

QIMRB_3

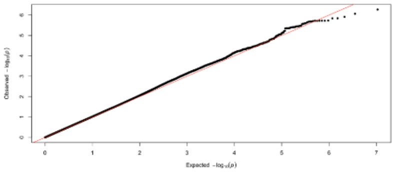

TRAILS

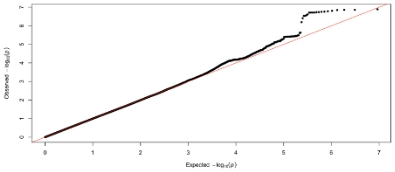

FinnTwin

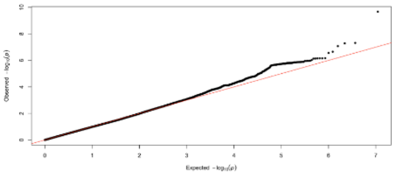

QIMRB_4

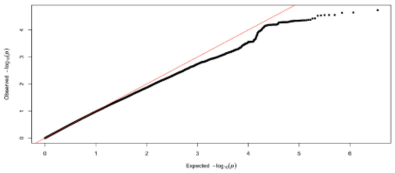

### 6. Manhattan and Q-Q plots discovery cohorts – combined

#### Manhattan plots

ADH

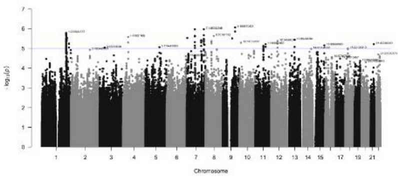

GSA

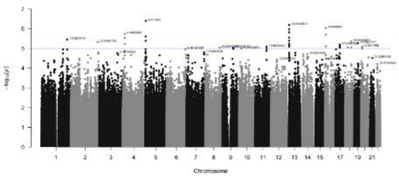

QIMRB_4

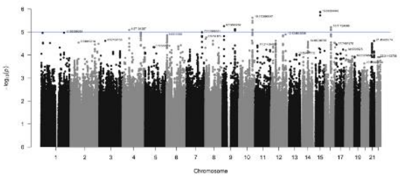

ALSPAC

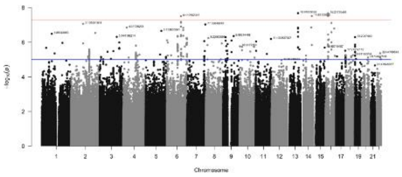

MCTFR

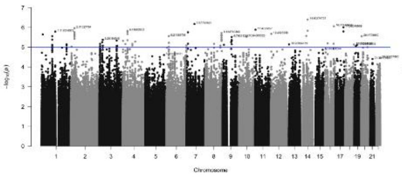

QIMRB_5

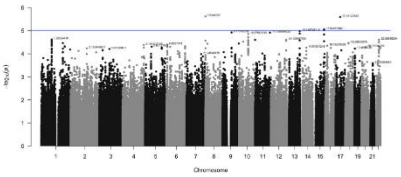

BIG

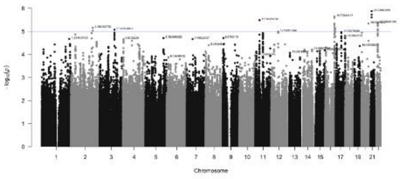

PAGES

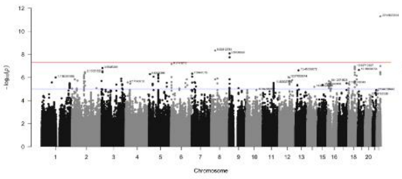

S42

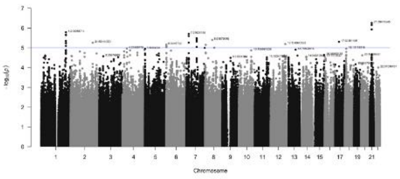

COGA

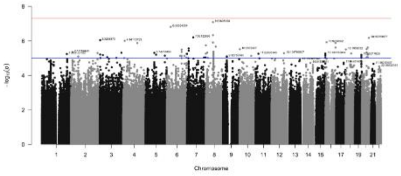

QIMRB_1

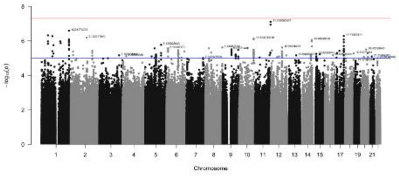

TEDS_1

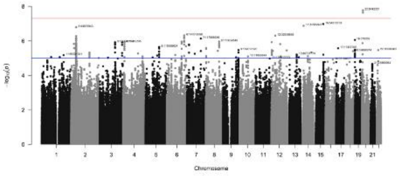

CoLaus

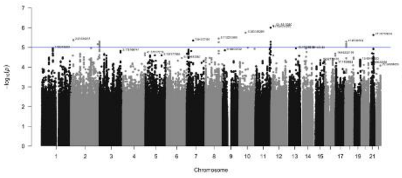

QIMRB_2

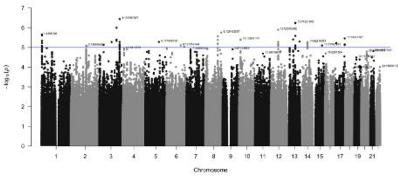

TEDS_2

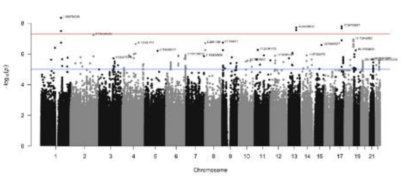

FinnTwin

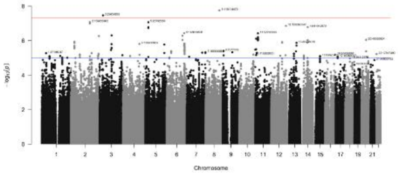

QIMRB_3

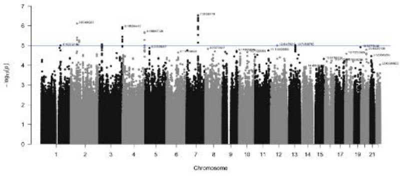

TRAILS

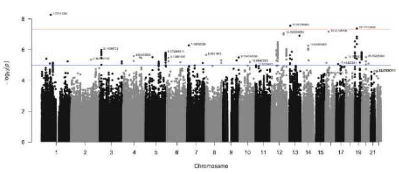

#### Quantile-quantile plots

ADH

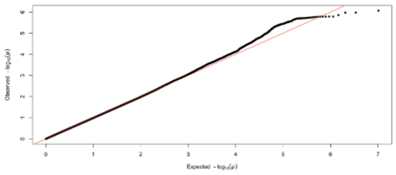

GSA

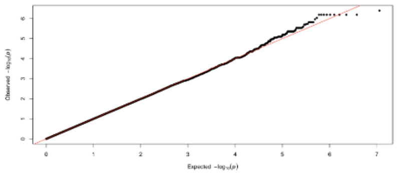

QIMRB_4

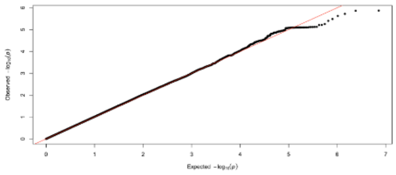

ALSPAC

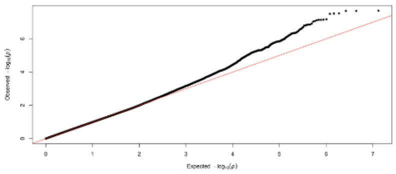

MCTFR

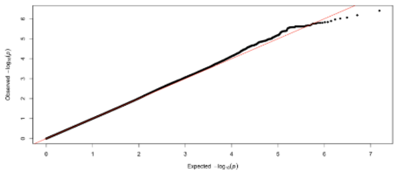

QIMRB_5

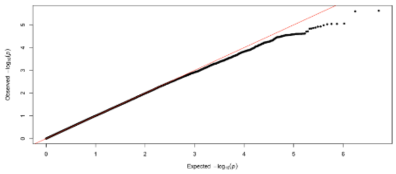

BIG

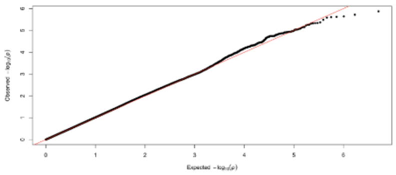

PAGES

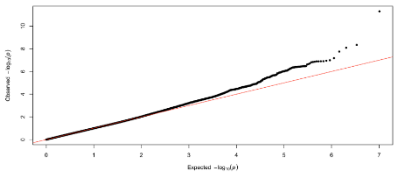

S42

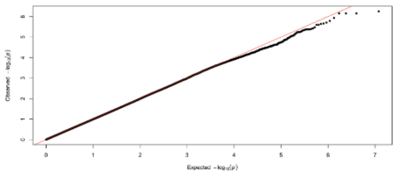

COGA

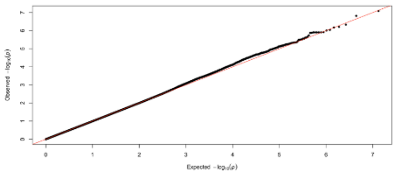

QIMRB_1

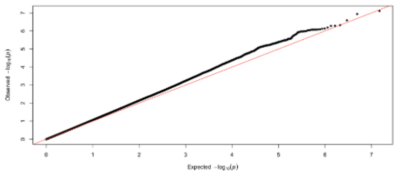

TEDS_1

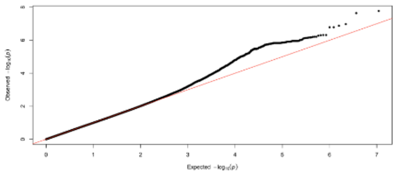

CoLaus

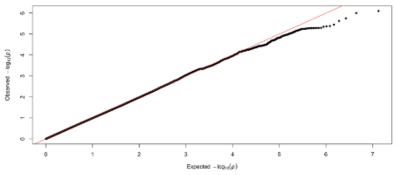

QIMRB_2

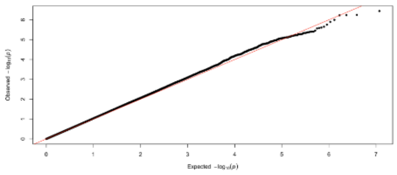

TEDS_2

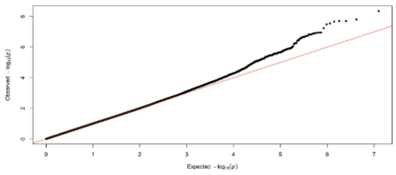

FinnTwin

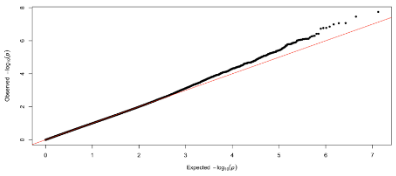

QIMRB_3

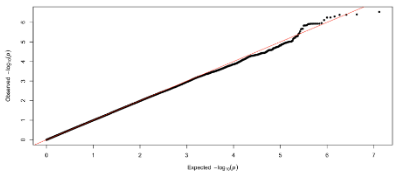

TRAILS

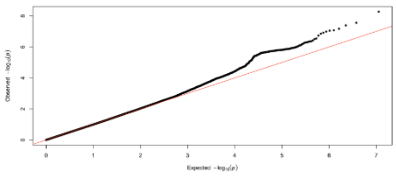

### 7. Sample and measurement description of independent cohorts

#### Dunedin Longitudinal Study

**Sample.** Participants in the second cohort were members of the Dunedin Multidisciplinary Health and Development Study, a longitudinal investigation of health and behavior in a birth cohort. Dunedin participants (N=1,037; 91% of eligible births; 52% male) were all individuals born between April 1972 and March 1973 in Dunedin, New Zealand, who were eligible on the basis of residence in the province and who participated in the first assessment at age 3. Full details about the sample are reported elsewhere (Poulton et al., 2015). The cohort represented the full range of socioeconomic status (SES) in the general population of New Zealand's South Island. On adult health, the cohort matches the New Zealand National Health and Nutrition Survey on key health indicators (e.g. body mass index, smoking, visits to the doctor).

Assessments were carried out at birth and ages 3, 5, 7, 9, 11, 13, 15, 18, 21, 26, 32, 38, most recently, 45 years, when 94% of the 997 participants still alive took part. At each assessment wave, participants are brought to the Dunedin research unit for a full day of interviews and examinations. These data are supplemented by searches of official records and by questionnaires that are mailed, as developmentally appropriate, to parents, teachers, and peers nominated by the participants themselves. The Otago Ethics Committee approved each phase of the study and informed consent was obtained from all participants.

**Genetic Data**. We used Illumina HumanOmni Express12 BeadChip arrays (Version 1.1; Illumina, Hayward, CA) to assay common single-nucleotide polymorphism (SNP) variation in the genomes of cohort members. We imputed additional SNPs using the IMPUTE2 software (Version 2.3.1; https://mathgen.stats.ox.ac.uk/impute/impute_v2.html; Howie *et al*, 2009) and the 1000 Genomes Phase 3 reference panel (1000 Genomes Project, 2012. Imputation was conducted on autosomal SNPs appearing in dbSNP (Version 140; http://www.ncbi .nlm.nih.gov/SNP/; (Sherry *et al*., 2001) that were "called" in more than 98% of the samples. Invariant SNPs were excluded. Pre-phasing and imputation were conducted using a 50-million-base-pair sliding window. The resulting genotype databases included genotyped SNPs and SNPs imputed with 90% probability of a specific genotype among the non-Maori members of the Dunedin cohort (n=918) and in Hardy-Weinberg equilibrium (p>0.01 for all). To address residual population stratification, we conducted a principal component analysis of our genome-wide SNP database using PLINK (Version 1.9; Chang *et al.* 2015).

**Measures.**

All Dunedin antisocial phenotypes have been previously published. We describe each measure and provide references to previous publications.

Parent and teacher reported child antisocial behavior. We assessed antisocial behavior at ages 5, 7, 9, and 11 years by using the Rutter Behaviour Checklist with mothers and teachers as the reporters (e.g., destroys property, fights, irritable, disobedient, tells lies, steals, bullies others.) We summed and standardized mothers' and teachers' reports of each of these measures to create a single cross-informant scale representing childhood antisocial behavior (Moffitt et al., 2011).

Conduct disorder was measured according to the criteria of the Diagnostic and Statistical Manual of Mental Disorders (DSM-IV), which identifies adolescents displaying a persistent pattern of behavior that violates the rights of others. A diagnosis of conduct disorder was made when we assessed the research participants at ages: 11, 13, 15. A 'lifetime' diagnosis was arrived at by establishing whether a Study member received the diagnosis at one or more of the three ages (Moffitt et al., 2011).

Criminal convictions. Information on officially recorded criminal offending was obtained by searching the central computer system of the New Zealand Police, which provides details of all convictions and sentences communicated to the New Zealand Police. Searches were completed following each assessment at ages 18, 21, 26, 32, 38, and 45. Official records of criminal conviction were available from 14 years of age onward, the age from which criminal conviction for all types of offenses was permissible. Criminal offending was recoded into a binary variable to reflect whether participants had ever been convicted or not.

Antisocial trajectories. We supplemented the analyses by studying developmental trajectories of parent, teacher, and self-reported antisocial behavior from childhood to adulthood. These trajectories have been developed and described in previous articles about antisocial behavior in the Dunedin cohort (Odgers et al., 2007; Odgers et al., 2008). Briefly, antisocial conduct problems were assessed at ages 7, 9, 11, 13, 15, 18, 21, and 26 years through scoring six key symptoms of DSM conduct disorder as being present or absent at each age according to reports from parents, teachers, and study members: physical fighting, bullying others, destroying property, telling lies, truancy, and stealing. Symptoms were adapted across the age span to ensure that the measures were developmentally appropriate (e.g., work absenteeism was substituted for truancy at older ages). Growth mixture modeling was used to identify subgroups of participants that followed unique trajectories in their antisocial behavior over time. A four-class model represented the best empirical fit to the data according to several indices of model fit and classification accuracy. Across gender, the majority of participants (50%) displayed low levels of antisocial behavior across time (the “always-low” group), 22% exhibited antisocial behavior only in childhood ("childhood-limited"), 19% were characterized by adolescent-limited antisocial behavior ("adolescent-limited"), and 9% displayed persistently high levels of antisocial behavior across the years ("life-course persistent").

Externalizing spectrum up to age 45 years. Participants' liability to externalizing psychopathology was measured at ages 18, 21, 26, 32, 38, and 45. Details are provided in a previous publication (Caspi et al., 2020). Briefly, we repeatedly assessed past-year symptoms of five disorders: DSM symptoms of alcohol dependence, cannabis dependence, hard drug dependence, and conduct disorder, as well as symptoms of tobacco dependence assessed with the Fagerström Test for Nicotine Dependence. Confirmatory factor analysis was used to derive a factor score (M=0, SD=1) indicating individuals' general risk for externalizing psychopathology from adolescence to midlife.

Partner violence. Intimate partner violence was assessed using the Dunedin Study Abuse Scales at ages 26, 32, 38, and 45 (Bourassa et al., 2020), a 33-item scale that used the items from the Conflict Tactics Scale–Revised (CTS-R), as well as additional items. The Physical Abuse scale included the nine physical violence items in the CTS-R, plus four additional items capturing other physically abusive behaviors (e.g., "Over the last year, did a partner ever push, grab, or shove you," and "hit, or try to hit you with something."). The Psychological Abuse scale consisted of two items from the CTS-R and 18 additional items capturing controlling, terrorizing, demeaning, and other psychologically abusive behaviors (e.g., "Over the last year, did a partner ever insult or shame you in front of others," and "Humiliate (or ridicule) you."). Study members were asked about their experience of partner violence victimization, as well as their perpetration of these 33 behaviors over the past 12 months in their romantic relationship. Study members reported whether each behavior occurred or not (0 not present,1 present), and these 66 values were summed to create an overall score for partner violence at each age, with higher scores representing greater partner violence. These scores were summed across the four phases to create an overall index of intimate partner violence.

#### Environmental Risk Longitudinal Study (E-Risk)

**Sample.** Participants were members of the Environmental Risk (E-Risk) Longitudinal Twin Study, which tracks the development of a birth cohort of 2,232 British participants. The sample was drawn from a larger birth register of twins born in England and Wales in 1994-1995. Full details about the sample are reported elsewhere (Moffitt et al., 2002). Briefly, the E-Risk sample was constructed in 1999-2000, when 1,116 families (93% of those eligible) with same-sex 5-year-old twins participated in home-visit assessments. This sample comprised 56% monozygotic (MZ) and 44% dizygotic (DZ) twin pairs; sex was evenly distributed within zygosity (49% male). Families were recruited to represent the UK population of families with newborns in the 1990s, on the basis of residential location throughout England and Wales and mother's age. Teenaged mothers with twins were over-selected to replace high-risk families who were selectively lost to the register through non-response. Older mothers having twins via assisted reproduction were under-selected to avoid an excess of well-educated older mothers. The study sample represents the full range of socioeconomic conditions in the UK, as reflected in the families' distribution on a neighborhood-level socioeconomic index (ACORN [A Classification of Residential Neighbourhoods], developed by CACI Inc. for commercial use): 25.6% of E-Risk families live in "wealthy achiever" neighborhoods compared to 25.3% nationwide; 5.3% vs. 11.6% live in "urban prosperity" neighborhoods; 29.6% vs. 26.9% live in "comfortably off" neighborhoods; 13.4% vs. 13.9% live in "moderate means" neighborhoods, and 26.1% vs. 20.7% live in “hard-pressed” neighborhoods. E-Risk underrepresents “urban prosperity” neighborhoods because such households are likely to be childless.

Home-visits assessments took place when participants were aged 5, 7, 10, 12 and, most recently, 18 years, when 93% of the participants took part. At ages 5, 7, 10, and 12 years, assessments were carried out with participants as well as their mothers (or primary caretakers); the home visit at age 18 included interviews only with participants. Each twin was assessed by a different interviewer. These data are supplemented by searches of official records and by questionnaires that are mailed, as developmentally appropriate, to teachers, and co-informants nominated by participants themselves. The Joint South London and Maudsley and the Institute of Psychiatry Research Ethics Committee approved each phase of the study. Parents gave informed consent and twins gave assent between 5-12 years and then informed consent at age 18.

**Genetic Data.** We used Illumina HumanOmni Express24 BeadChip arrays (Version 1.1; Illumina, Hayward, CA) to assay common single-nucleotide polymorphism (SNP) variation in the genomes of cohort members. We imputed additional SNPs using the IMPUTE2 software (Version 2.3.1; https://mathgen.stats.ox.ac.uk/ impute/impute_v2.html; Howie *et al.*, 2009) and the 1000 Genomes Phase 3 reference panel (Genomes Project Consortium, 2012). Imputation was conducted on autosomal SNPs appearing in dbSNP (Version 140; http://www. ncbi.nlm.nih.gov/SNP/; Sherry *et al.*, 2001) that were “called” in more than 98% of the samples. Invariant SNPs were excluded. The E-Risk cohort contains MZ twins, who are genetically identical; we therefore empirically measured genotypes of one randomly selected twin per pair and assigned these data to their MZ co-twin. We directly measured genotypes of both members of dizygotic twin pairs. Prephasing and imputation were conducted using a 50-million-base-pair sliding window. The resulting genotype databases included genotyped SNPs and SNPs imputed with 90% probability of a specific genotype among the European descent members of the E-Risk cohort (N=1,999 participants in 1,011 families).

To address residual population stratification, we conducted a principal component analysis of our genome-wide SNP database using PLINK (Version 1.9; Chang *et al.* 2015). One twin was selected at random from each family for principal component analysis. SNP loadings for principal components were applied to co-twin genetic data to compute principal component values for the full sample.

**Measures.**

All E-Risk antisocial phenotypes have been previously published. We describe each measure and provide references to previous publications.

Parent and teacher reported child antisocial behavior. We assessed childhood antisocial behavior at ages 5, 7, 10, and 12 years by using Achenbach's Child Behavior Checklist in interviews with mothers and the Teacher Report Form by mail with teachers (Wertz et al., 2016). We summed and standardized mothers' and teachers' reports of each of these measures to create a single cross-informant scale representing childhood antisocial behavior.

Conduct disorder was measured according to the criteria of the Diagnostic and Statistical Manual of Mental Disorders (DSM-IV). A diagnosis of conduct disorder was made on the basis of symptoms reported by mothers and teachers when the participants were ages 5, 7, 10, and 12 years. A 'lifetime' diagnosis was arrived at by establishing whether a Study member received the diagnosis at one or more of the four ages (Wertz et al., 2018).

Criminal conviction. Official records of participants' criminal offending were obtained through UK Police National Computer record searches conducted in cooperation with the UK Ministry of Justice. Records include complete histories of cautions and convictions beginning at age 10, the age of criminal responsibility. Our data are complete through age 22. Criminal offending was coded as a binary variable to reflect whether participants had been cautioned or convicted.

Externalizing spectrum. Participants' liability to externalizing psychopathology was measured at age 18. Details are provided in a previous publication (Schaefer et al., 2018). Briefly, we assessed past-year symptoms of five disorders: DSM symptoms of alcohol dependence, cannabis dependence, conduct disorder, ADHD, as well as symptoms of tobacco dependence assessed with the Fagerström Test for Nicotine Dependence. Confirmatory factor analysis was used to derive a factor score (M=0, SD=1) indicating individuals' general risk for externalizing psychopathology in adolescence.

#### Quebec Longitudinal Study of Child Development (QLSCD)

**Sample.** Participants in the third cohort were part of the Quebec Longitudinal Study of Child Development, an ongoing longitudinal study of a representative sample of children, starting at the age of 5 months, who were sampled from the Quebec birth registry to be representative of the population of infants born in 1997/1998 in the province of Québec, Canada. Full details about the sample are reported in Orri et al. (2020). More than 2,000 families participated in the study when the infant was aged 5 months in 1998 (T1, n=2,120). These children were assessed at home, and their parents interviewed yearly (six waves overall) before school entry. The children all entered grade school in 2003 (Kindergarten), and were assessed regularly in the winter-spring period of each assessment year, starting in 2004 (Kindergarten). Data have been collected annually or every other year up to age to 21 years. The cohort currently includes 1245 participants. At each assessment wave, information on a wide range of child and family characteristics, environmental, biological (e.g., hair cortisol, genetic, epigenetic), and administrative data was gathered. The QLSCD protocol was initially approved by the Health Quebec ethics committee.

**Measures.** Anti-social behaviors were self-assessed through the Mental Health and Social Inadaptation Assessment for Adolescents (MIA) at age 19. The MIA is a self-report instrument for quantifying through a dimensional approach the mental health symptoms and psychosocial adaptation problems based on the DSM-5. The instrument includes a variety of subscales pertaining to both internalizing disorders and externalizing disorders, including Conduct Disorder under which self-ratings refer to (1) lying and cheating, (2) stealing and breaking the law, (3) breaking rules, and (4) vandalism. Scores were derived by summing the item. For more information on the content of the MIA, its reliability and reliability, see Côté et al., (2018).

#### Quebec Newborn Twin Study (QNTS)

**Sample.** Participants in the fourth cohort were part of the Quebec Newborn Twin Study, an ongoing prospective longitudinal follow-up of a birth cohort of twins born between 1995 and 1998 in the greater Montreal area, Québec, Canada. The goal of QNTS is to document individual differences in the cognitive, behavioral, and social-emotional aspects of developmental health across childhood, their early genetic and environmental determinants, as well as their putative role in later social-emotional adjustment, school, health, and occupational outcomes. A total of 662 families participated in the first wave of assessments (i.e., at age 6 months on average). These twin children, and their parents, were then assessed longitudinally in the lab and at home during preschool (5 waves), and then in grade school (9 waves), and up to age 21 (ongoing). Full details about the sample and follow-up are reported here: Boivin et al., (2013, 2019). Participants were assessed longitudinally on a broad range of characteristics, including anti-social behaviors, through multi-informant, multi-method assessments of child outcomes. QNTS also entailed extended and detailed multilevel assessments of proximal (e.g., parenting behaviors) and distal (e.g., family income) features of the twins' environment across development. The QNTS offers unique features for the study of cognitive, behavioral, and social-emotional development during childhood, and for examining the contribution of the early developmental trajectories and contexts to the child's later adjustment and long-term health benefits. The QNTS also shares many of its features (measures and time of assessments) with the Québec Longitudinal Study of Child Development. Ethical approval of study protocol was obtained from the institutional review committees of Université Laval and Hôpital Sainte-Justine, as well as the councils of the participating schools.

**Measures.** Anti-social behaviours were assessed at 20 years old via the Achenbach System of Empirically Based Assessment (ASEBA) questionnaire (Achenbach, 2015). The ASEBA is a 20-item self-report questionnaire assessing adaptive and maladaptive functioning.

Teachers also rated a variety of antisocial behaviors in primary school using a variant of the Child Social Behavior Questionnaire, which centers on a variety of externalizing (e.g., hyperactivity, impulsivity, inattention, and aggression) and internalizing (e.g., anxiety, shyness, social withdrawal, depression symptoms) behaviour problems from 6 to 12 years.

##### Genetic Data for QLSC and QNTS

Genetic Data. Genotype data were collected from blood or saliva samples from children of European descent. Genotyping was performed at Genome Quebec, Montreal, Canada using Illumina PsychArray-24 v1.3 Beadchip. We imputed additional SNPs using SHAPEIT v2 (r837) (Delaneau et al., 2013), IMPUTE2 v2.3.2 (Bryan N. Howie et al., 2009), and the 1000 Genomes Phase 3 reference panel. After imputation, variants with a MAF <1%, an HWE test p <1×10-6, and an INFO metric <0.8 were removed. The resulting genotype databases included 8,407,807 genotyped and imputed SNPs.

To address residual population stratification, we conducted a principal component analysis of our genome-wide SNP database using PLINK v1.90b6.7 for QLSCD and PLINK v1.90b5.2 for QNTS. The first ten components were saved for later use as covariates in the main analyses to control for genetic stratification.

#### Philadelphia Neurodevelopmental Cohort (PNC)

Sample. The Philadelphia Neurodevelopmental Cohort (PNC) is a large sample of genotyped children, adolescents, and young adults (ages 8-21) in the United States who have completed a variety of clinical and cognitive phenotypic measures (Satterthwaite et al., 2016). All relevant data were downloaded via the Database for Genotypes and Phenotypes (dbGaP Study Accession: phs000607.v3.p2). As requested by the original study investigators, we refer the reader to previous publications for additional information on the collection of genetic (Glessner et al., 2010) and clinical phenotypic data (Calkins et al., 2014, 2015). In order to minimize the threat of population stratification, we limited analyses in the present study to individuals who (i) self-reported non-Hispanic European ancestry and (ii) were not identified as ancestral outliers per genetic principal component analysis. Furthermore, participants missing responses on 10% or more of items were excluded from statistical analyses. The relevant PNC data is briefly described below.

Genetic data. Genotypes were prepared and processed in accordance with recommendations for chip-based genomic data (Anderson et al., 2010; Turner et al., 2011). We excluded samples from statistical analysis on the basis of poor call rate (< 98%) and inconsistent self-reported sex and biological sex. We also pruned the cohort for relatedness, removing one sample at random from each kinship pair (KING coefficient > .0442). We then excluded variants from analysis if more than 2% of genotype data was missing. As filtering on minor allele frequency or Hardy-Weinberg equilibrium (HWE) has been shown to have a detrimental effect on imputation quality (Roshyara et al., 2014), these filters were applied after phasing and imputation.

Untyped variants were imputed on the Michigan Imputation Server (https://imputationserver.sph.umich.edu). Genotypes were phased with Eagle v2.3 (Loh et al., 2016) and imputed with Minimac3 1.0.13 (Das et al., 2016) while using the 1000 Genomes Project Phase 3 v5 (1000 Genomes Project Consortium, 2015). To ensure markers were of high quality, we applied several post-imputation quality control thresholds, filtering (i) SNPs with a MAF < 0.01, (ii) imputation quality score < 0.60, or (iii) HWE P value < 1e-6 were excluded from statistical analyses. Finally, we restricted variants to biallelic HapMap3 SNPs when constructing polygenic scores.

As previously noted, the present analyses were limited to participants who self-reported non-Hispanic European descent. However, to further account for cryptic relatedness and population stratification, we used flashPCA2 (Abraham, Qiu and Inouye, 2017) to compute the top five principal components of ancestry. These scores were also used to identify ancestral outliers (i.e., participants with substantial levels of admixture), and included as covariates in all statistical analyses.

Measures. Symptoms of oppositional defiant disorder and conduct disorder were assessed using the GOASSESS (Satterthwaite et al., 2016), a computerized assessment battery designed to comprehensively and systematically evaluate psychopathology in pediatric samples. The GOASSESS can be considered to be an abbreviated and modified form of the epidemiological version of the Kiddie Schedule for Affective Disorders and Schizophrenia (K-SADS; Merikangas et al., 2009), which assesses lifetime occurrence of symptoms across most major domains of psychopathology. In the present study, symptoms were modelled in a latent variable framework. Age, sex, and principal components of ancestry were included as covariates

### 8. Polygenic risk scoring results in Dunedin and E-risk

#### The Dunedin cohort

The PRS for ASB was associated with higher scores on the Rutter Behaviour Checklist. For the Rutter scale the increase in R2 with the addition of PRS to the model ($\Delta$ ($R^2$ PRS+sex) - ($R^2$ sex)), incremental R2, was .02 in the most optimal model (P-value threshold of .01), indicating that the PRS could explain up to 2.0 % of the variance in antisocial behavior (Supplementary Table 9).

The ordinary least squares (OLS) regression demonstrates - using p-value thresholds of .05 and .1 - that a higher PRS for ASB (*P* = .019 and *P* = .021 respectively, OLS Regression) was associated with higher symptom counts on Conduct Disorder in children. Moreover, a higher PRS for ASB was linked to more symptom counts on both Conduct Disorder and Antisocial Personality Disorder reported by mothers (*P* = .042, OLS Regression) and fathers (*P* = .033 and *P* = .04 respectively). Higher PRS for ASB was also associated with the number of adult court convictions up to age 45, with elevated Incident Rate Ratio's (*P* = .01 for P-value thresholds of .0001). The increased association with PRS becomes even more pronounced when also juvenile convictions are being considered (OR= 1.35, p < .001 for several P-value thresholds).

For externalizing behavior, assessed longitudinally from age 18 to age 45, we found an incremental R2 of .9% (for P-value thresholds of .05 and .1). Examining the antisocial trajectories we found that the life course persistent antisocial behavior (LCP) group had a significantly (p = .032 and p =.049, for P-value thresholds .05 and .1 respectively) higher ASB PRS score as compared to the group that did not display antisocial behavior (No; Supplementary Figure 2). Lastly, we did not find evidence for association between our PRS for ASB and exposure to partner violence (p > .05 for all P-value thresholds).

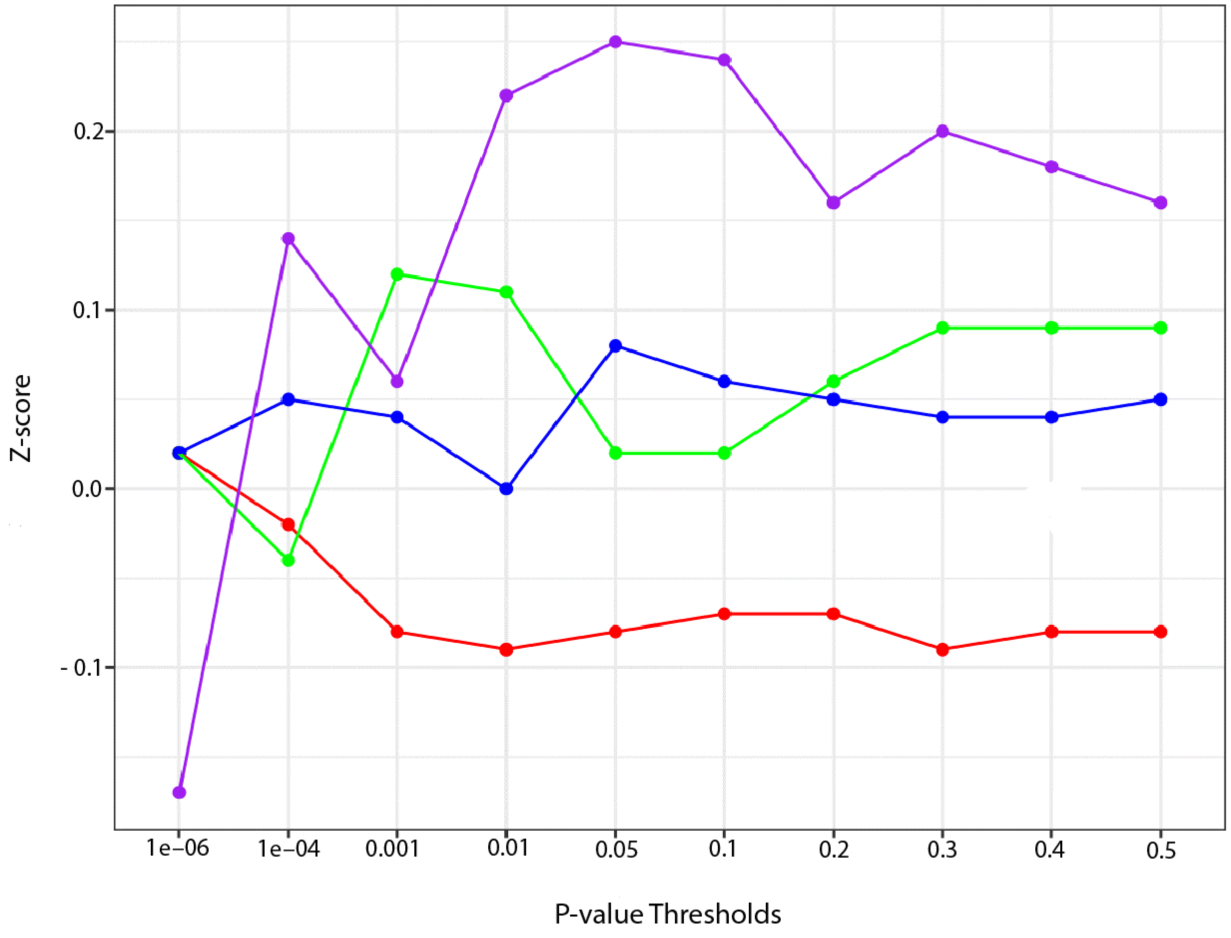

Supplementary Figure 2. The highest genetic risk of broad ASB was associated with individuals following the life-course persistent antisocial trajectory (shown in purple) followed by the childhood-limited antisocial behavior (shown in green) and adolescent-onset antisocial behavior (shown in blue) trajectories, the lowest genetic risk was associated with the low ASB trajectory (shown in red).

#### Environmental Risk Longitudinal Study (E-Risk)

When examining parent and teacher reported antisocial behavior checklists up to age 12, we found that the total score was associated with a higher PRS for ASB ($P < .001$, for all P-value thresholds below .1). Comparison of the full model (polygenic score + covariates) with a covariates-only model, indicated that the PRS could explain up to 3.7% of the variance in childhood psychopathology. Next, we demonstrate that a higher PRS for ASB is associated with higher symptom counts on CD as reported by mothers and teachers ($P < .001$, for all P-value thresholds below .1, OLS Regression). This positive association was also found for any conduct disorder diagnosis by mothers and teachers ($P < .001$, for all P-value thresholds below .1).

In addition, when considering the number of criminal offences up to age 22, we found that the PRS was linked to elevated Incident Rate Ratio's ($P < .01$ for several P-value thresholds). Moreover, the twin difference analyses showed that the PRS could significantly predict differences in the externalizing problems scale (aggressive and delinquent behavior) between 433 dizygotic twin pairs (r=.135, $P = .005$, for P-value threshold .2). Similarly, looking at externalizing behavior at age 18, we found significant associations ($P < .001$, for all P-value thresholds below .01) and the PRS could explain up to 3.7% of the variance in externalizing behavior (Supplementary Table 10). The twin difference analyses did not reveal a significant correlation between twin pair differences in the 3-factor model of externalizing behavior and differences in PRS ($P > .05$).
